# Select EZH2 inhibitors enhance the viral mimicry effects of DNMT inhibition through a mechanism involving calcium-calcineurin-NFAT signaling

**DOI:** 10.1101/2023.06.09.544393

**Authors:** Alison A. Chomiak, Rochelle L. Tiedemann, Yanqing Liu, Xiangqian Kong, Ying Cui, Kate Thurlow, Evan M. Cornett, Michael J. Topper, Stephen B. Baylin, Scott B. Rothbart

## Abstract

DNA methyltransferase (DNMT) inhibitors are FDA-approved for various hematological malignancies but have limited efficacy in solid tumors. DNA hypomethylation with these drugs is associated with elevated lysine 27 tri-methylation on histone H3 (H3K27me3). We hypothesized that this EZH2-dependent repressive mark limits the full potential of DNMT inhibition. Here, we show in cell line and tumoroid models of colorectal cancer, that low-dose DNMT inhibition sensitizes cells to selective EZH2 inhibitors that have limited single agent toxicity, and that EZH2 inhibition enhances DNMT inhibitor-driven molecular and therapeutic effects. Through integrative epigenomic analyses, we reveal that DNMT inhibition induces H3K27me3 accumulation at genomic regions poised with EZH2. Unexpectedly, combined treatment alters the epigenome landscape to promote transcriptional upregulation of the calcium-calcineurin-NFAT signaling pathway. Blocking this pathway limits the transcriptional activating effects of the drug combination, including expression of transposable elements and innate immune response genes within a viral defense pathway. Consistently, we demonstrate positive correlations between DNMT inhibitor- and innate immune response-associated transcription profiles and calcium signal activation in primary human colon cancer specimens. Collectively, our study demonstrates that compensatory EZH2 activity following DNA hypomethylation presents a barrier to the therapeutic action of DNMT inhibition in colon cancer, reveals a new application of EZH2 inhibitors beyond cancers associated with PRC2 hyperactivity, and links calcium-calcineurin-NFAT signaling to epigenetic therapy-induced viral mimicry.

**Highlights:**

- Select EZH2 inhibitors enhance the transcriptional activating and antiproliferative effects of DNA hypomethylating agents in colon cancer cells.
- The mechanism involves blockade of H3K27me3 accumulation in regions of the genome poised for PRC2 activity.
- DNMT inhibitor + EZH2 inhibitor treatment transcriptionally upregulates calcium-calcineurin-NFAT signaling, and this pathway is necessary for complete induction of viral mimicry and innate immune response pathways.
- The therapeutic utility of EZH2 inhibitors may be extended beyond cancers with PRC2 hyperactivity in combination regimens with DNMT inhibitors.

## Introduction

Abnormal DNA methylation patterning, and its associations with altered gene expression, is recognized as an enabling hallmark of human cancers (1). These cancer-specific changes include global DNA hypomethylation coupled with focal DNA hypermethylation at CpG-rich promoters (i.e., CpG islands; CGIs), which serves as a major mechanism of tumor suppressor gene (TSG) silencing (2–5). As such, DNA methyltransferase (DNMT) inhibitors, including the nucleoside analogs 5- azacytidine (AZA) and 5-aza-2’-deoxycytidine (DAC, decitabine), have been deployed as an epigenetic cancer management strategy with the goal of reversing DNA methylation-mediated transcriptional silencing. DNMT inhibitors also induce a “viral mimicry” response triggered by de-repression of endogenous retroviruses (ERVs) and other transposable elements (TEs) that stimulate dsRNA- and dsDNA-dependent expression of innate immune and inflammasome signaling pathways (6–9). These data suggest that DNA hypomethylation therapy has the potential to enhance the efficacy of immune checkpoint therapy, as has been demonstrated in pre-clinical models (10–12).

AZA and DAC have demonstrated clinical successes with FDA approvals for the treatment of acute myeloid leukemia (AML) and myelodysplastic syndrome (MDS), and their use correlates with DNA hypomethylation and transcriptional activation of TSGs and TEs in these hematological malignancies (13–17). While DNMT inhibitor therapy has provided superior survival benefit for some patients with AML or MDS (14), only a subset of these patients have had a complete response and others develop resistance to therapy (16,18–24). Notably, the short half-life of AZA and DAC, and their dependence on DNA replication for drug activity, diminishes their accumulation at doses that induce favorable transcriptional outcomes. This is particularly problematic in solid tumors, where DNMT inhibitor efficacy has been limited (25–27). Unfortunately, these efficacy barriers are unlikely to be addressed with modified dosing, as high dose DNMT inhibition incurs DNA damage and cytotoxicity, thereby reducing DNMT inhibitor target activity (25,28–30). In contrast, repeated exposure to low dose DNMT inhibitors allows these nucleoside analogs to exert stronger and more sustained epigenetic effects (17,25,31–37). As the high potential achieved in pre-clinical studies has yet to be observed in the many ongoing clinical trials of DNMT inhibition, an unmet need exists for better combinatorial therapeutic approaches that enhance the epigenetic effects of DNMT inhibitors within patient-tolerable doses (13).

EZH2 is a histone methyltransferase subunit of Polycomb Repressive Complex 2 (PRC2) that catalyzes all three states of lysine 27 methylation on histone H3 (H3K27me) (38,39). PRC2 activity is essential in differentiation and development, where H3K27me3, a histone post-translational modification (PTM) associated with facultative heterochromatin and transcriptional repression, co-occurs at select promoters with H3K4me3, a histone PTM associated with active transcription (40,41). This “bivalent state” is most commonly found at unmethylated CGI promoters of repressed lineage commitment genes in stem cells, allowing for rapid gene activation or continued repression by the “poised” PRC2 complex, depending on the cellular differentiation cues received (42–45). In cancer, bivalent genes repressed by H3K27me3 early in development are often stably silenced through the acquisition of CGI promoter DNA hypermethylation (3,42–51). Targeting this pathologic DNA hypermethylation through genetic or chemical disruption of DNMTs results in a re-emergence of the stem-like chromatin state associated with PRC2 activity that reinforces transcriptional silencing (42,43,45,47,52).

We hypothesize that compensatory silencing through H3K27me3 may, in part, explain the barrier to robust and complete clinical responses to DNMT inhibitor therapy, providing rationale for combining DNMT inhibitors with EZH2 inhibitors. In support of this hypothesis, recent studies have shown that DNA hypomethylation, induced genetically or as a consequence of chemotherapeutic and metabolic perturbation, sensitizes cancer cells to EZH2 inhibition (53–55). Moreover, combination anti- neoplastic/therapeutic effects on multiple cancer cell lines and tumor xenografts have been reported following combined treatment with DNMT and EZH2 inhibitors (11,56–59). As clinical applications of EZH2 inhibition have primarily focused on cancers addicted to PRC2 activity resulting from activating mutations in EZH2 or loss of function mutations in subunits of the SWI/SNF chromatin remodeling complex (60–64), combining EZH2 inhibitors with DNMT inhibitors presents an opportunity to expand the utility of these epigenetic agents for cancer therapy.

In this study, we sought to dissect the relationship between DNA methylation and PRC2 activity as repressive transcriptional regulatory mechanisms in colorectal cancer (CRC) and to evaluate the molecular effects of blocking these two nodes of epigenetic signaling as a potential cancer management strategy. While CRC, a solid tumor type that is one of the leading causes of global cancer deaths (65), has been well-characterized for its abnormal DNA methylation patterns, it has limited single-agent sensitivity to EZH2 and DNMT inhibition. Comparing a panel of commonly used and/or clinically tested EZH2 inhibitors revealed that EPZ6438 (tazemetostat, TAZ), the most clinically advanced EZH2 inhibitor with FDA approvals for epithelioid sarcoma and follicular lymphoma (62,63), enhanced the transcriptional activating and antiproliferative effects of DNMT inhibition across multiple CRC cell lines. While TAZ was the most potent of the EZH2 inhibitors tested, this surprisingly did not correlate with single agent toxicity. The limited toxicity of TAZ in these cell types enables transcriptional synergy with DNMT inhibitors, whose mechanism of action requires rounds of DNA replication in the absence of DNMT activity. Indeed, transcriptome-wide analyses revealed that EZH2 inhibition enhanced known DNMT inhibitor-responsive pathways, including upregulation of pathways associated with the innate immune response and viral mimicry (6,8) and downregulation of *MYC* and MYC target genes (66). Notably, the calcium-calcineurin-NFAT signaling pathway, which is traditionally associated with T-cell activation, also emerged as a top enriched pathway following acute combination treatment. Converging on this observation, integration of transcriptomics with ChIP-seq and DNA methylation analyses revealed that regions of the genome that accumulate H3K27me3 in response to DNA hypomethylation are positioned over regulatory elements for AP1, Jun/FOS, and NFAT target genes that are poised with EZH2. Blocking intracellular calcium signal transduction prevented DNMT and EZH2 inhibitor-induced transcriptional upregulation of the calcium signaling pathway and limited transcriptional activating effects of this drug combination towards TEs, innate immune, and inflammation-associated pathways, revealing an unexpected connection between these signaling axes in the molecular response to epigenetic therapy. In agreement with these findings, high expression of innate immune response genes in tumor biopsies from CRC patients correlated with expression of the upregulated calcium signaling genes identified in our study.

Altogether, we show that CRC cells that are insensitive to single agent EZH2 inhibition become sensitive as compensatory H3K27me3-mediated silencing emerges upon DNMT inhibitor treatment, enriching new transcriptional pathways as a consequence of the epigenetic effects. Epigenomic and transcriptomic analyses converged on the calcium-calcineurin-NFAT signaling pathway as a place where this epigenetic shift coincides with a transcriptional response. These new insights into the molecular mechanisms of epigenetic crosstalk suggest a rational drug combination that could enhance therapeutic responses to clinically applied DNMT inhibitor epigenetic therapies while also directing EZH2 inhibitors in new clinical directions.

## Results

### Select EZH2 inhibitors synergize with DNMT inhibitors to reactivate a TSG silenced by DNA hypermethylation

The appreciation that certain cancer types are addicted to PRC2 activity has resulted in the development of numerous EZH2 inhibitors that are routinely used in the laboratory setting and clinical trials (67–69). We first considered the extent to which many of the most commonly used EZH2 inhibitors augment the transcriptional activating effects of DAC by querying effects of combined EZH2 and DNMT inhibition on the expression of the TSG *SFRP1.* This gene is epigenetically silenced by promoter CGI hypermethylation in colon cancer and was shown to acquire H3K27me3 following DNA hypomethylation (52,70). To facilitate this, we used CRISPR/Cas9 to insert a NanoLuciferase (NLuc) cassette into exon 2 of the endogenous *SFRP1* locus of HCT116 cells (Figure 1A), a colon cancer cell line that also shows an EZH2-dependent global increase in H3K27me3 following DAC treatment (**Figure S1A**). Of note, this reporter was engineered in an HCT116 DNMT1 hypomorphic (MT1) cell line characterized by ∼20% reduction in global DNA methylation to enhance the sensitivity to DNA hypomethylating agents (71). NLuc signal, a proxy for *SFRP1* expression, was measurable following siRNA-mediated knockdown of DNMT1 and in a dose- and time-dependent manner following treatment with DAC (Figure 1B).

**Figure 1.**
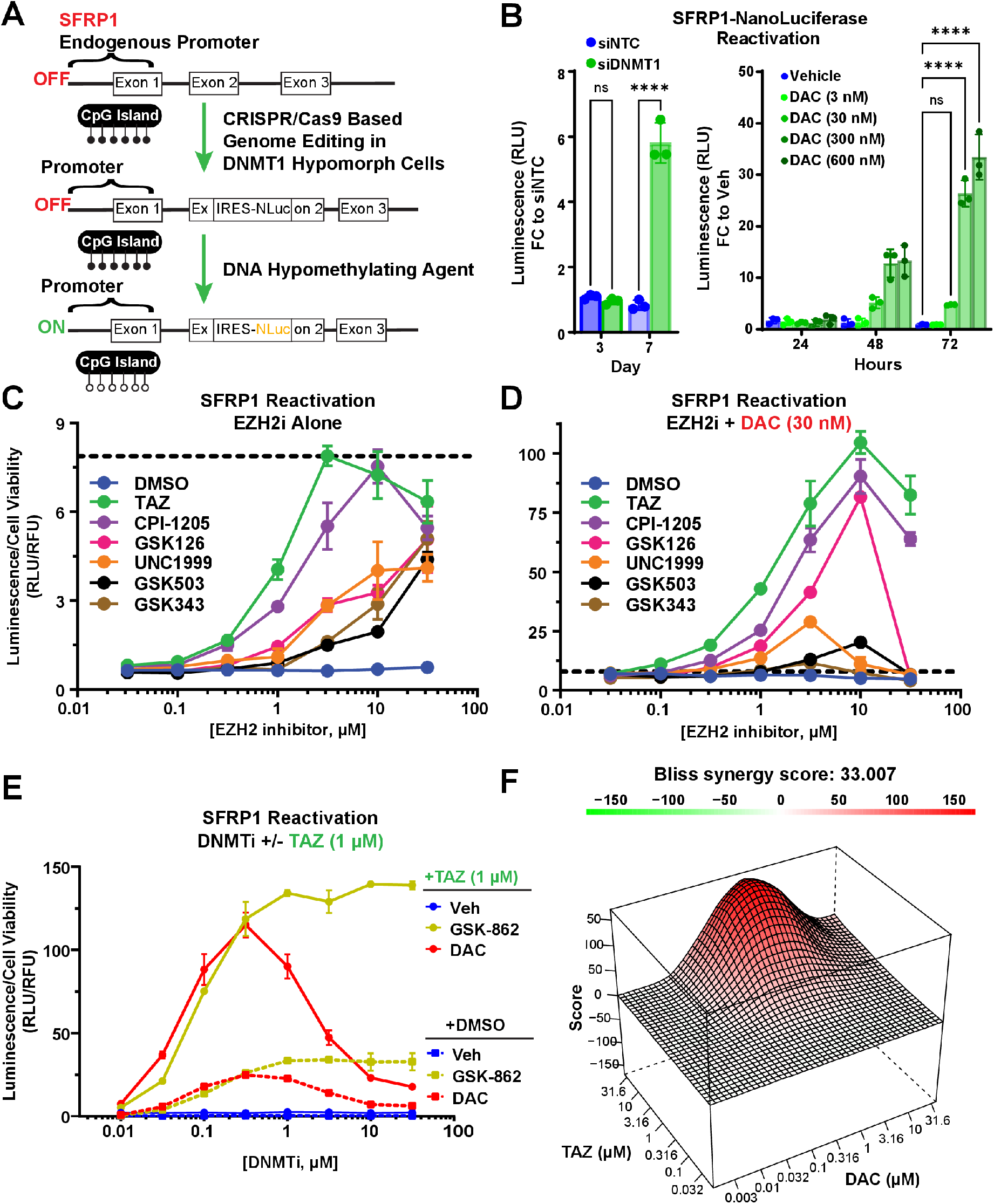
Blocking PRC2 activity with select EZH2 inhibitors enhances the transcriptional activating effects of DNA hypomethylating agents. **A)** Schematic of the approach to engineer an in- cell NanoLuciferase (NLuc) reporter assay to measure expression from an endogenous allele of the tumor suppressor gene *SFRP1* in HCT116 cells with hypomorphic DNMT1. **B)** Reactivation of the silenced *SFRP1* locus from the *SFRP1-*NLuc reporter assay following DNA hypomethylation via siRNA- mediated DNMT1 knockdown (left) or treatment with DAC (right). Relative Luminescence Units (RLU) are shown as fold change over control at the first timepoint. NTC, siRNA non-targeting control. Data are presented as mean +/- SD of technical triplicates. ****p<0.0001. **C-D)** NLuc reporter activity measurements following 72-hour treatment with the indicated EZH2 inhibitors **(C)** alone or **(D)** in combination with a fixed concentration of DAC. Error bars are presented as mean +/- SD of technical triplicates. Dotted black lines denote scaling difference between panels **C** and **D**. **E)** NLuc reporter activity measurements following 72-hour treatment with the DNMT inhibitors DAC or GSK3484862 alone (dashed lines) or in combination with a fixed concentration of TAZ (solid lines). Error bars are presented as mean +/- SD of technical triplicates. **F)** Surface plot and Bliss synergy score using SynergyFinder2.0 to analyze combination DAC plus TAZ dose response curves derived from the *SFRP1*-NLuc reporter assay after 72-hour treatment.

Using this NLuc reporter assay, we screened a panel of S-adenosyl-L-methionine (SAM)- competitive small molecule inhibitors of EZH2 that included TAZ, CPI-1205, EPZ005687, EPZ011989, GSK126, GSK343, GSK503, and UNC1999 (**Figure S1B**) (61,72–80). TAZ and CPI-1205, the most clinically advanced among these compounds, and EPZ011989 (a derivative of TAZ), induced *SFRP1* expression and subsequent NLuc reporter activity following an acute 72-hour treatment over a broad range of concentrations, whereas other EZH2 inhibitors showed marginal induction of *SFRP1* expression as single agents (Figure 1C and **S1C-D**). Combining these EZH2 inhibitors with a fixed low dose of DAC (30 nM) that had limited single agent activity in the NLuc assay and minimal to moderate effects on DNA methylation loss (Figure 1B and **S1E**) enhanced the maximum measured luminescence by more than ten-fold over the same range of concentrations compared to single agent EZH2 inhibition (Figure 1D and **S1C-D**). Again, this observation was strongest for TAZ and CPI-1205 over the other EZH2 inhibitors.

We next considered the extent to which TAZ, the best performing EZH2 inhibitor, augments the transcriptional activating effects of DNMT inhibition by combining it at a fixed dose with either DAC or the non-nucleoside DNMT1 inhibitor GSK3484862 over a range of concentrations. These two molecules induce comparable dose-dependent loss of DNA methylation across the CGI promoter of *SFRP1* in wild-type HCT116 (**Figure S1F**). However, the maximum NLuc reporter assay signal plateaued and remained elevated at high doses of GSK-862, whereas DAC showed a bell curve of activity with a maximum NLuc signal at 300 nM (Figure 1E). This difference is related to the induction of toxicity by DAC at high doses, which is suggested, by comparison to GSK-862, to be DNA hypomethylation-independent. Strikingly, the addition of TAZ amplified DNMT inhibitor-induced upregulation of the *SFRP1* NLuc reporter by more than 10-fold (Figure 1E). Pairwise dose-response titrations of TAZ and DAC confirmed the synergistic interaction in the NLuc reporter assay (Figure 1F and **S1G**). From this, Bliss synergy scores of 33.007 and 131.984 for treatment durations of three or six days, respectively, were calculated, which far exceeded the threshold of 10 indicative of a synergistic relationship (81) (Figure 1F and **S1G**). The selective ability for TAZ to synergize with DAC was validated by qRT-PCR of *SFRP1* expression in wild-type and MT1 HCT116 (**Figure S1H**). Collectively, these data show that select EZH2 inhibitors synergize with DNMT inhibitors to elevate expression of a TSG known to be silenced by DNA hypermethylation.

### Single agent EZH2 inhibitor potency and therapeutic efficacy do not correlate

We assumed that the superior activity of TAZ and CPI-1205 to reactivate *SFRP1* was related to their enhanced potency as EZH2 inhibitors. Indeed, western blot analysis of HCT116 cell lysates showed that both TAZ and CPI-1205 dose-dependently reduced global levels of H3K27me3 and did so at lower concentrations than four other EZH2 inhibitors after 72 hours of treatment (Figure 2A and **Figure S2A-B**) despite similar *in vitro* IC50s for PRC2 inhibition (**Figure S2C**). Unexpectedly, potency did not predict cellular toxicity. Dose-response cell viability assays showed both MT1 and wild-type HCT116 were viable over a much broader range of concentrations when treated with TAZ or CPI-1205 compared to GSK126, GSK503, GSK343, or UNC1999 (Figure 2B and **Figure S2D**). These findings of toxic but less potent EZH2 inhibition, despite similar *in vitro* IC50s against PRC2, suggest off-target activity separate from their ability to inhibit EZH2. In comparison, the ability of TAZ and CPI-1205 to strongly inhibit PRC2 with limited toxicity is ideal for maximizing the epigenetic effects of DNMT inhibitors that require cell replication and DNA synthesis over a prolonged course of treatment.

**Figure 2.**
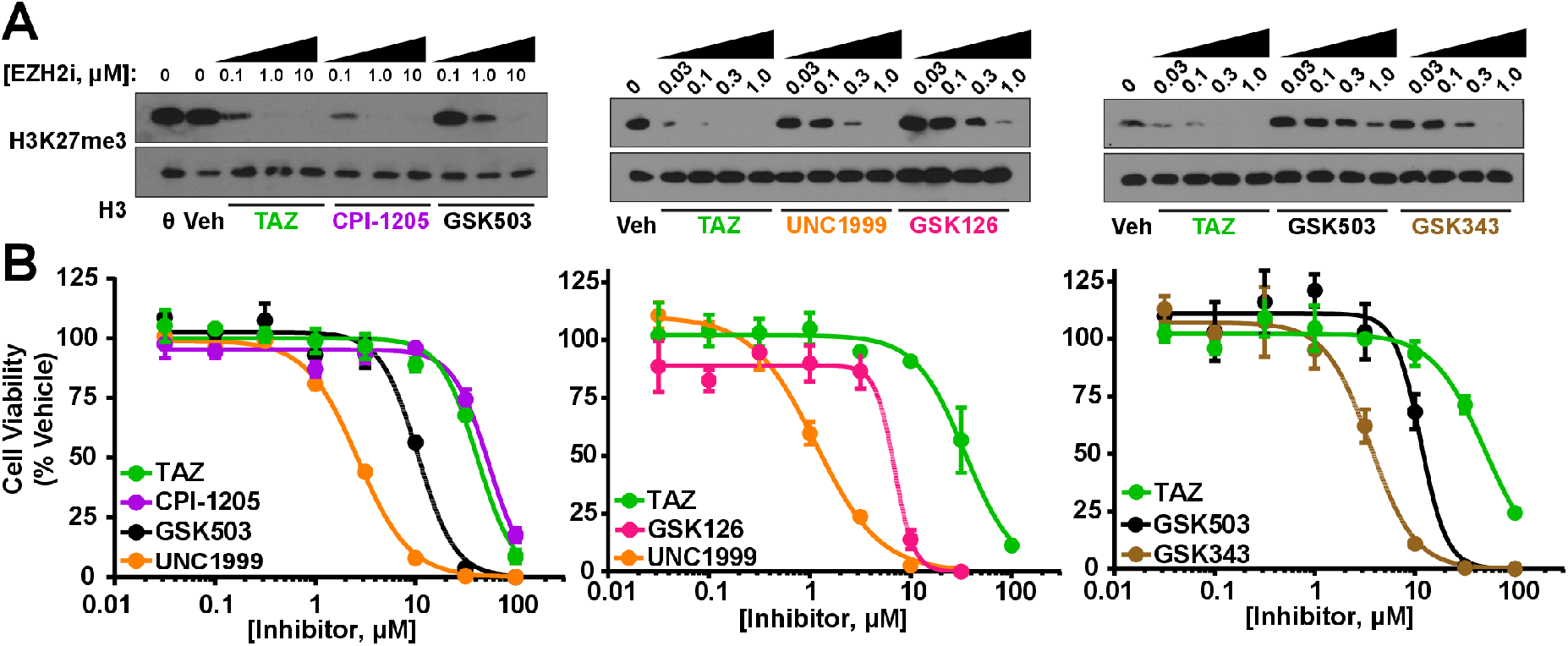
Single agent EZH2 inhibitor potency and therapeutic efficacy do not correlate in HCT116. **A)** Western blot analysis of H3K27me3 from wild-type HCT116 cells following 72-hour exposure to vehicle (DMSO, % equivalent) or EZH2 inhibitors at the indicated concentrations. **B)** Dose response viability curves (CellTiter-Fluor normalized to vehicle) for *SFRP1*-NLuc Reporter/DNMT1 hypomorph HCT116 cells exposed to the indicated EZH2 inhibitors for 72 hours. Data are presented as mean +/-SD of technical triplicates.

### DNMT and EZH2 inhibitors have combination therapeutic efficacy in cell line and tumoroid models of colon cancer

We next sought to determine whether DNMT inhibition could sensitize colon cancer cells to EZH2 inhibition. We first compared a potent but non-toxic EZH2 inhibitor (TAZ) to a less potent but more toxic EZH2 inhibitor (GSK343) in combination with DAC. Indeed, when exposed to drugs for 7 days, DAC plus TAZ showed combined efficacy at reducing cell viability (Figure 3A). Conversely, the strong single agent toxicity of GSK343 reduced cell viability to such an extent acutely (3 days) and over time (7 days) that it prevented any therapeutic cooperation with DAC (Figure 3A).

**Figure 3.**
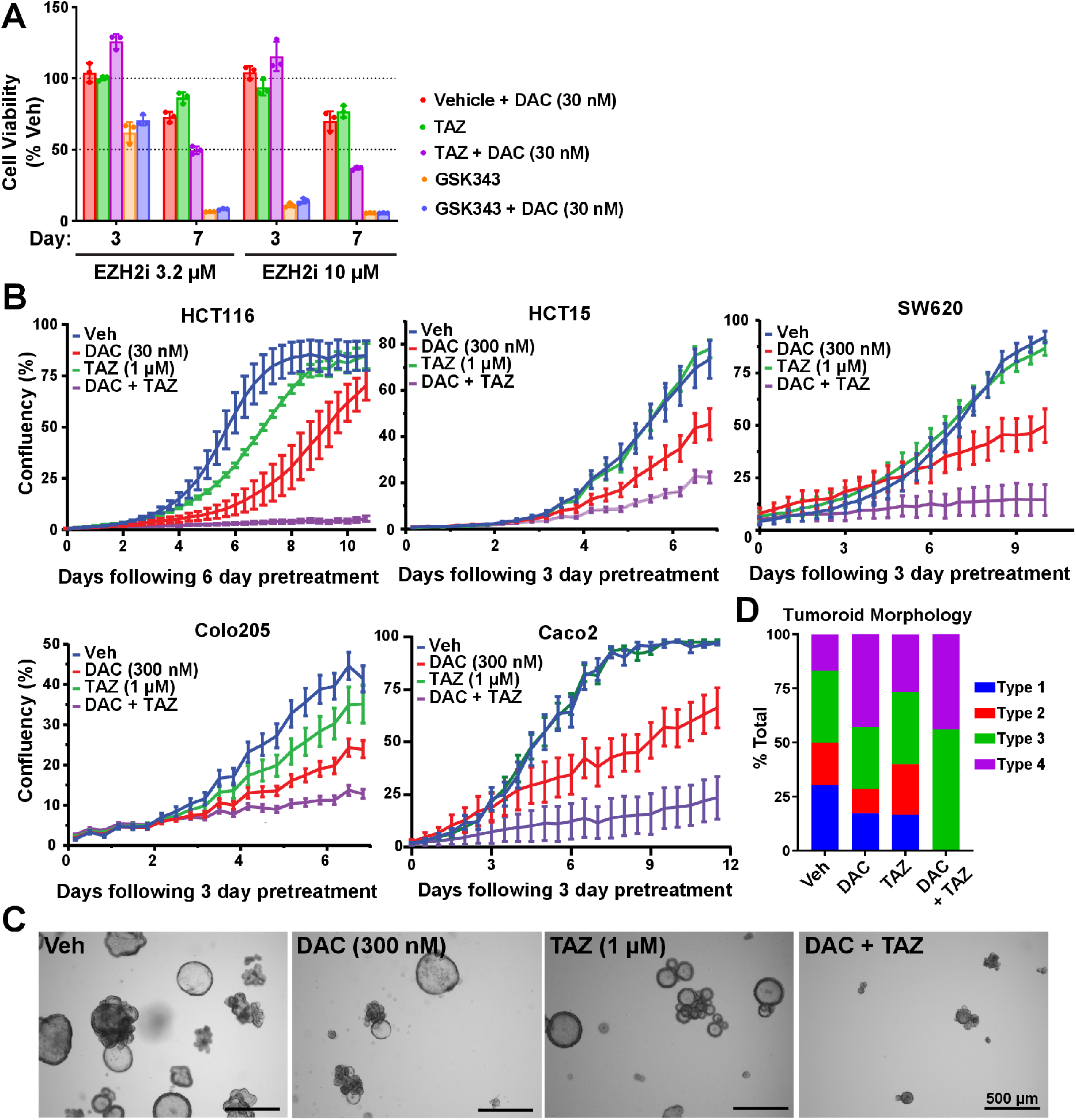
DAC and TAZ synergize to reduce colon cancer cell and tumoroid proliferation. **A)** CellTiter-Fluor viability assays for the HCT116 *SFRP1-*NLuc reporter/DNMT1 hypomorph cell line after 3 or 7 days of exposure to the indicated EZH2 inhibitor with or without DAC. Data are presented as mean +/-SD of technical triplicates. **B)** Confluency measurements from Incucyte imaging of the indicated cell lines. Cells were split and drugs were refreshed every 72 hours for the indicated number of days prior to plating (“pretreatment”) and were re-treated and re-plated again at Day 0 of Incucyte measurements. Data are presented as mean +/- SEM of technical replicates averaged from 12 images (HCT116, HCT15, and COLO205) or 9 images (SW620 and CACO2) per timepoint and treatment. **C)** Representative brightfield images of *ex vivo* tumoroids derived from an APC^Min+^ mouse intestinal adenoma and treated in culture for 21 days. Vehicle or inhibitors were refreshed every 72-96 hours, and tumoroids were split every 7-9 days. **D)** Quantification of tumoroid growth/shape based on observed phenotypes (Types I and II representing normal and viable and Types III and IV representing least viable/healthy and loss of cyst-like morphology as shown in **Figure S3H**.) Data are presented as % of total tumoroids evaluated, n=35-66 organoids/treatment group. See also **Figure S3**.

We extended this observation across a panel of colon cancer cell lines representative of the major genetic and epigenetic subtypes of this disease (82). Again, as a single agent, TAZ had limited anti-proliferative effects in these cell lines at doses that were sufficient to induce loss of H3K27me3 (Figure 3B and **S3A-B**) and that far exceeded the IC50s of this drug in sensitive cell types (74,83,84). However, when combined with doses of DAC that showed limited single agent toxicity in culture and reflect the degree of DNA methylation loss achieved in clinical trials (**Figure S1C**) refs (23), strong combination antiproliferative effects were observed across the CRC cell line panel. These combination antiproliferative effects were most pronounced following multiple low-dose treatments over a period of 6 to 9 days (Figure 3B and **S3D**), despite evidence of DNA and histone hypomethylation as early as 3 days at these concentrations (**Figure S3A, C**), or after single agent exposures at elevated concentrations (Figure 3A and **S3E-G**).

Similar observations were made following *ex vivo* treatment of small intestine-derived tumoroids from APC^Min/+^ mice (Figure 3C). While tumoroids treated with TAZ appeared morphologically similar to vehicle-treated tumoroids, tumoroids treated with DAC showed a reduction in size, arrested growth, and loss of the standard cyst-like, hollowed spherical morphology with a clearly defined lumen (Figure 3D and **S3H**). (85–88). Combination treatment dramatically enhanced this phenotype. Collectively, these data show that TAZ and DAC have combined therapeutic efficacy in a temporal- and concentration-dependent manner, limiting cell proliferation in multiple colon cancer cell line and tumoroid models that are insensitive to single agent EZH2 inhibition.

### DNMT inhibition induces H3K27me3 at genomic regions poised with EZH2

The combined molecular and therapeutic efficacy of DNMT and EZH2 inhibition, despite limited effects with EZH2 inhibition alone, suggested that a PRC2 dependency emerges following hypomethylation of the genome. To test this hypothesis, we next performed integrative epigenomic analysis of quantitative distributions of DNA methylation and H3K27me3 in response to these drug treatments. To query DNA methylation distributions, we first utilized the Illumina MethylationEPIC bead array (EPIC array), a platform that interrogates the DNA methylation status of ∼850,000 unique CpGs across the genome with a biased distribution for probes covering promoters, gene bodies, enhancers, and intergenic space. Single-agent TAZ treatment had no effect on DNA methylation distribution, and the combination of TAZ with a low dose of DAC (30 nM, DAC30) did not further deplete DNA methylation relative to DAC treatment alone (**Figure S1E**, **S4A-B**). Moreover, DNA methylation loss induced by both single-agent DAC and the combination with TAZ occurred at the same CpG loci (**Figure S4A**) and to the same degree of DNA hypomethylation (**Figure S4B**, average Δβ-value ≅ −0.14 for both DAC30 and Combo). Finally, these conserved DNA hypomethylation events between single-agent and combination DAC treatment occurred primarily across gene promoters and distal regulatory space (both intergenic and genic) (**Figure S4C**).

Using our recently developed sans spike-in quantitative ChIP-seq method (siQ-ChIP) (89,90), we next quantified changes in H3K27me3 following drug exposure (**Figure S4D-G**). TAZ treatment, alone and in combination with DAC, significantly reduced H3K27me3 levels across the genome following 3 days of treatment (Figure 4A). To query where H3K27me3 loss primarily occurs in the genome after TAZ treatment, we built custom ChromHMM annotations for parental HCT116 cells (**Figure S4H-I**). As expected, the predominant regions of the genome that lost H3K27me3 with either single agent or combination TAZ treatment occurred in annotated Polycomb Repressive regions (ReprPC1, ReprPC2, EnhBiv) that had abundant H3K27me3 signal in vehicle-treated cells (Figure 4C, left panel). In addition to ChromHMM annotations, we considered new genomic studies that place H3K27me3 at crucial transition regions of the genome. H3K27me3 is now appreciated to demarcate an intermediate “I” compartment of the genome that sits between the active ‘A’ compartment and inactive ‘B’ compartment (91,92). Additionally, these ‘I’ compartments are enriched for the transition between early and late replication timing (93) (**Figure S4J**). Using publicly available 16-phase Repli-seq data for HCT116 (94), we asked how H3K27me3 changes following treatment in the context of replication timing phases. Consistent with the notion that H3K27me3 is enriched at transitional regions, loss of H3K27me3 was enriched for the phases of replication timing that denote a transition from early replication to late replication (Figure 4D, left panel). Collectively, our analysis demonstrates that TAZ, in both the presence and absence of DAC, induces significant loss of H3K27me3 at genomic regions known to be regulated by PRC2.

**Figure 4.**
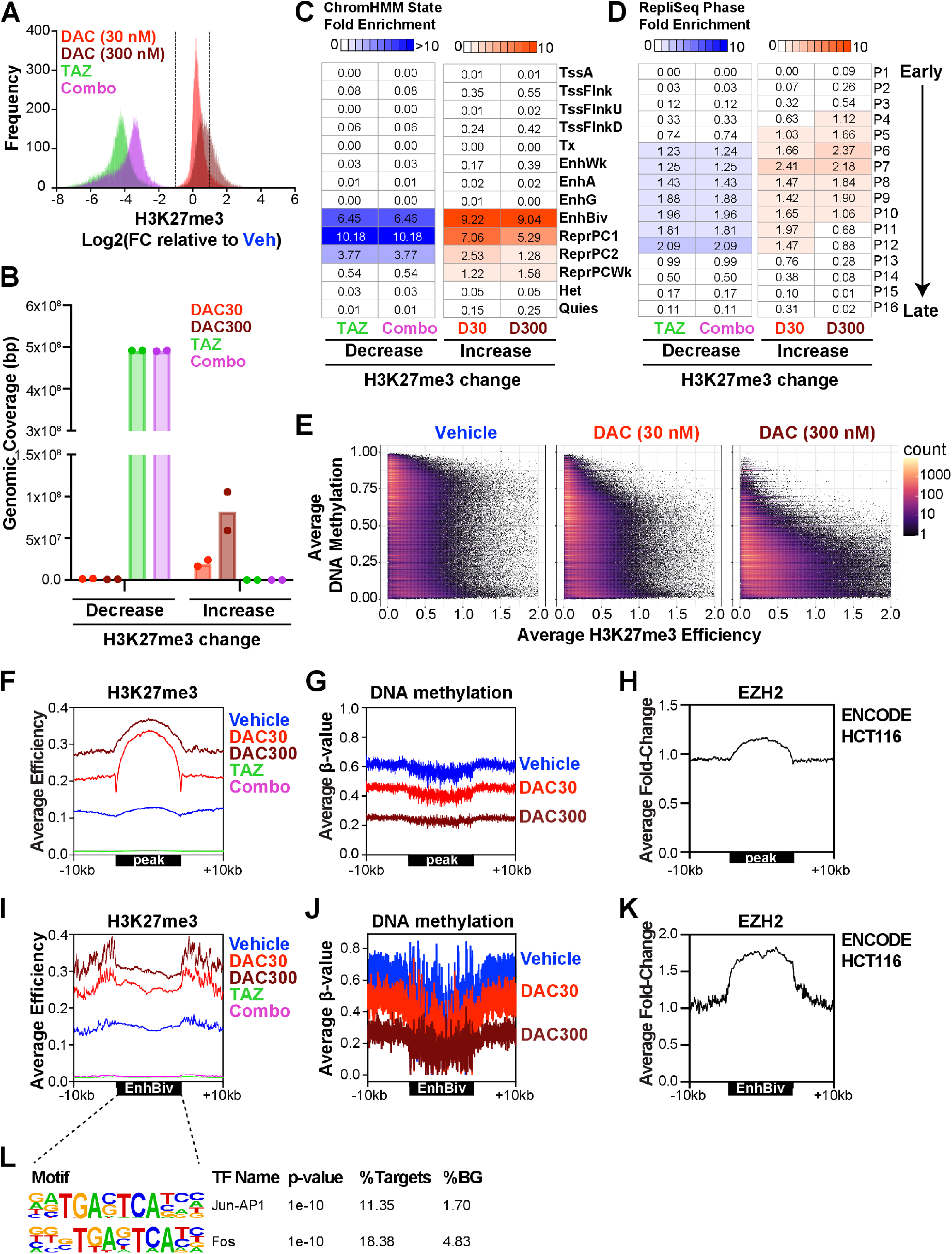
Increased levels of H3K27me3 are associated with DAC-induced DNA hypomethylation at genomic regions poised with EZH2. **A)** Histograms of Log_2_(FC) in H3K27me3 efficiency relative to Vehicle treated samples. Combo is 1 µM TAZ plus 30 nM DAC. Log_2_(FC) is averaged across biological duplicates. Dotted lines indicate the threshold for H3K27me3 signal considered significantly different from Vehicle (|Log_2_FC| ≥ 1). **B)** Genomic coverage (in bp) of significantly altered regions of H3K27me3 efficiency (|Log_2_FC| ≥ 1) relative to Vehicle. **C-D)** Fold enrichment overlap analysis of lost (TAZ/Combo) and gained (DAC 30 nM/DAC 300 nM) H3K27me3 efficiency regions relative to Vehicle (|Log_2_FC| ≥ 1) in relation to **C)** ChromHMM states and **D)** replication timing (RepliSeq) phases from HCT116. D30 and D300, 30 nM DAC and 300 nM DAC. **E)** Density scatterplots of average H3K27me3 efficiency (measured by siQ-ChIP-seq) versus average DNA methylation (measured by EM-seq of siQ-ChIP-seq fragments) for the indicated treatment conditions following 72-hour exposure in HCT116. Average values are plotted from biological duplicates across 100 bp bins. **F-H)** Average profiles for **F)** H3K27me3 efficiency, **G)** DNA methylation, and **H)** EZH2 occupancy at conserved genomic regions that show increased H3K27me3 levels following 30 nM DAC treatment (n=3,223 peaks). **I-K)** Average profiles for **I)** H3K27me3 efficiency, **J)** DNA methylation, and **K)** EZH2 occupancy at conserved Bivalent Enhancers (EnhBiv) that show increased H3K27me3 following 30 nM DAC treatment (n=185 enhancer regions). **L)** HOMER transcription factor motif enrichment analysis of EnhBiv regions that show increased H3K27me3 with 30 nM DAC treatment. %Targets, % of given regions with indicated motif; %BG, % of the queried representative background with indicated motif. See also **Figure S4**.

From prior work, we hypothesized that H3K27me3 would replace DNA methylation as a repressive epigenetic signal when genomes are hypomethylated (42,43,45,47,52). Consistent with this hypothesis, western blot analysis showed that H3K27me3 levels globally increased following DNMT inhibitor treatment, and this increase was captured in a dose-dependent manner with siQ-ChIP (Figure 4A-B and **S1A**). ChromHMM and Repli-seq enrichment analysis demonstrated that increases in H3K27me3 occurred at Polycomb Repressive regulated regions (EnhBiv, ReprPC1, ReprPC2, ReprPCWk) and intermediate replication timing phases, respectively (Figure 4C-D, right panels).

Importantly, while single agent TAZ and combination treatments did not induce significant losses of H3K27me3 in “weak” Polycomb regions (ReprPCWk) (Figure 4C, left panel), the increases in H3K27me3 following DAC treatment did occur in these regions (Figure 4C, right panel).

By global DNA methylation analysis on the EPIC array, low-dose DAC in single agent and combination treatments effectively reduced DNA methylation levels by approximately 15% (**Figure S1E**, **S4B**). The prevailing view in the field is that H3K27me3 and DNA methylation genomic distributions are mutually exclusive. However, experiments coupling H3K27me3 ChIP-seq with bisulfite sequencing (ChIP-BS-seq) of immunoprecipitated fragments identified genomic regions where these two epigenetic modifications co-occur and showed that this dual repression epigenetic signature is more prevalent in cancer cells (58,95,96). As the EPIC array provides limited coverage of Polycomb Repressive genomic regions, we employed a similar technique by coupling siQ-ChIP to Enzymatic Methyl (EM)-seq (ChIP-EM-seq) to generate high-coverage maps of the quantitative distributions of DNA methylation and H3K27me3 in cells exposed to DNMT inhibition, which allowed us to define the relationship between these epigenetic modifications on the same chromatin fragments (**Figure S4D-G**). Consistent with DAC-induced global accumulation of H3K27me3 (**Figure S1D**), DNMT inhibitor single agent treatment effectively decreased DNA methylation while increasing H3K27me3 in a dose-dependent manner (Figure 4E). Importantly, as the levels of H3K27me3 increased, the average DNA methylation levels decreased. However, while H3K27me3 and DNA methylation do appear to largely anti-correlate with each other, many instances exist where these modifications do co-exist in the genome (**Figure S4G**).

Next, we focused on these identified genomic regions that consistently demonstrated increased H3K27me3 following low-dose DAC treatment and analyzed the gains in H3K27me3 averaged over the start and stop coordinates of these regions (Figure 4F, **S4K**). Notably, these peaks start with low levels of H3K27me3 in vehicle treated controls and significantly increased H3K27me3 in a dose-dependent manner following DAC treatment. Importantly, these regions of the genome are most prone to undergo an ‘epigenetic switch,’ replacing DNA methylation with H3K27me3 following DAC treatment (Figure 4F-G). ENCODE ChIP-seq data from HCT116 showed enrichment of EZH2 across these regions (Figure 4H, **S4L**), suggesting that PRC2 is ‘poised’ in these regions for H3K27me3 deposition when the genome is hypomethylated. Given the localization of hypomethylated differentially methylated regions (DMRs) at intronic, genic, and distal intergenic regions (**Figure S4C**), we next determined that bivalent enhancers (regulatory elements dually marked by H3K4me1 and H3K27me3) follow the ‘epigenetic switch’ pattern (Figure 4I-J) and these enhancers also appear to be poised with EZH2 to mediate H3K27me3 accumulation (Figure 4K). Notably, HOMER motif analysis revealed a significant enrichment for AP-1 (Jun/Fos) binding motifs at these bivalent enhancers undergoing an ‘epigenetic switch’ (Figure 4L). Collectively, our epigenomic analyses suggest that combination treatment effectively induces loss of DNA methylation, blocks deposition of H3K27me3, and opens up enhancers for AP-1 transcription factor accessibility.

### EZH2 inhibition enhances transcriptional responses associated with DNMT inhibition after prolonged treatment

Although significant alterations to the global epigenome were observed as early as 3 days post-treatment (**Figure S1A, E**), antiproliferative effects of combined DNMT and EZH2 inhibition were most prominent with extended treatment. Therefore, we next sought to analyze the global transcriptional response to the DAC plus TAZ combination following both prolonged (6 days) and acute (3 days) treatments. Single agent DAC and TAZ drug treatments upregulated hundreds of protein-coding (PC) genes on their own (Figure 5A-D). Notably, combination treatments further enhanced the transcriptional upregulation of these single agent-responsive genes (**Figure S5A-D**) along with stimulating the expression of over 600 additional PC genes following both prolonged (Figure 5A-B) and acute treatments (Figure 5C-D). A comparison of genes upregulated at both acute and prolonged treatment intervals showed that many genes upregulated at day 6 were already expressed by day 3 of treatment for both single agent and combination treatment arms (**Figure S5E**). Prolonged treatment revealed that TEs were also significantly upregulated with single agent treatments, and to a much greater extent with the combination (Figure 5A). Notably, TE expression following acute treatment was unique to combination treatment and increased nearly 10-fold following prolonged drug exposure (Figure 5C), indicating that re-expression of transposable elements requires sustained, prolonged treatment with the DNMT and EZH2 inhibitor combination when DNMT inhibitors are used at a low dose. This suggests induction of viral mimicry, and an associated innate immune response, may be acutely initiated with the drug combination, but that a robust induction requires prolonged drug exposure.

**Figure 5.**
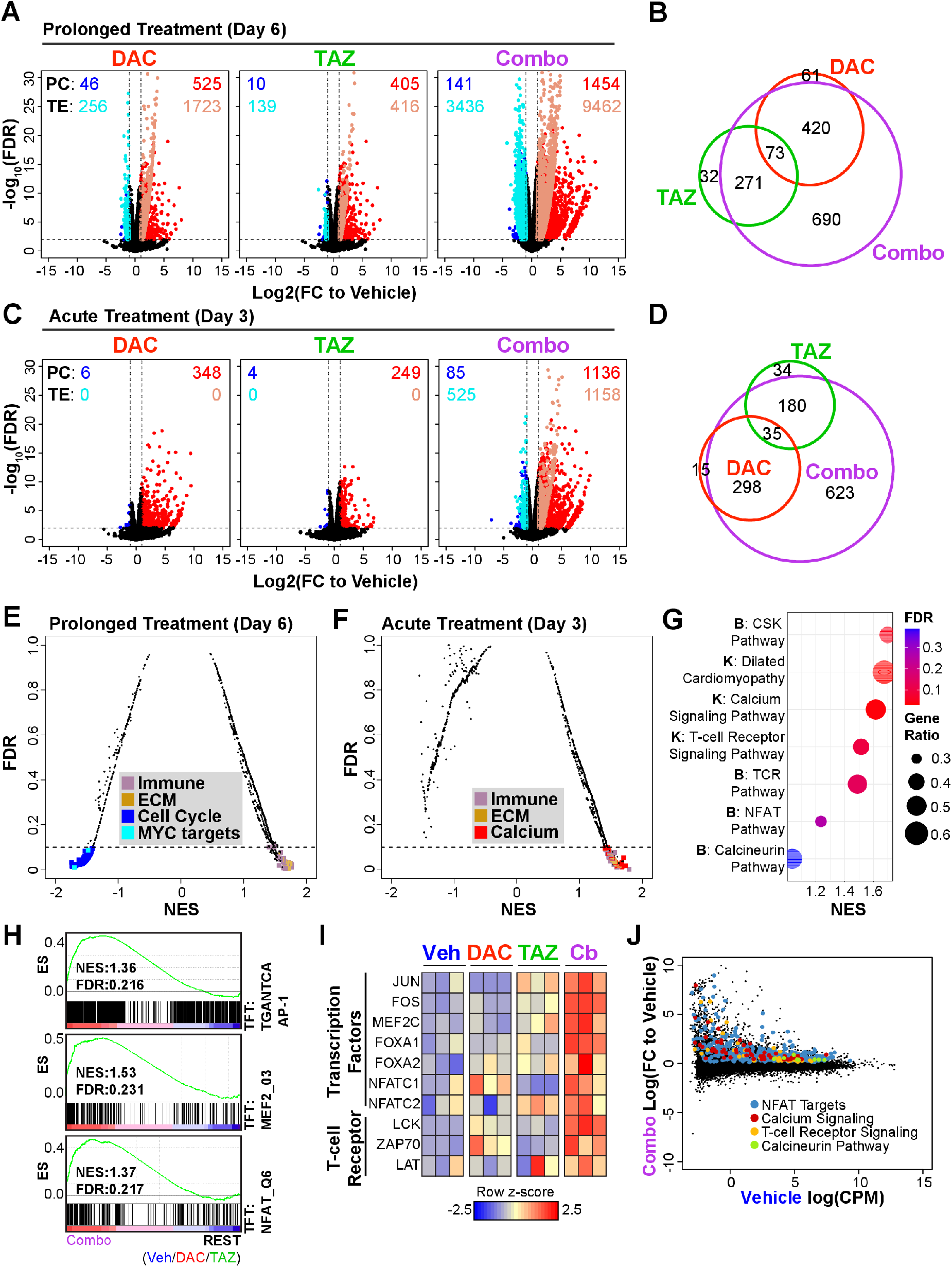
TAZ enhances transcriptional responses associated with DNMT inhibition. **A)** Volcano plots for protein-coding (PC) genes and transposable element (TE) expression from total RNA-seq of indicated inhibitor treatments (DAC, 30 nM; TAZ, 1 µM) relative to Vehicle in HCT116 following 6 days of treatment. Cells were split and re-treated with inhibitors every 72 hours. Data points are plotted as averages from three biological replicates. PC and TE transcripts that meet a significance threshold of Log_2_(FC) ≥ 1.0 and FDR ≤ 0.01 are colored points. Black points are PC transcripts that do not meet threshold conditions. Non-significant TE transcripts were excluded from the panel for simplicity. **B)** Venn diagram of overlapping PC genes significantly upregulated by single agent or combination inhibitor treatment from **A**. **C)** Volcano plots for PC genes and TE expression from total RNA-seq among indicated inhibitor treatments (DAC, 30 nM; TAZ, 1 µM) relative to Vehicle (DMSO/PBS) in HCT116 cells after single treatment for 72 hours. Data points are plotted as averages from three biological replicates. PC and TE transcripts that meet a significance threshold of Log_2_(FC) ≥ 1.0 and FDR ≤ 0.01 are colored. Black points are PC transcripts that do not meet threshold conditions. **D)** Venn diagram of overlapping PC genes significantly upregulated by single agent or combination inhibitor treatment from panel **C**. **E)** Gene Set Enrichment Analysis (GSEA) summary for HALLMARK, REACTOME, KEGG, and BIOCARTA gene sets in Combo treatment versus REST (Veh/DAC/TAZ) after six days of treatment. Gene sets that meet a significance threshold of FDR (False Discovery Rate) ≤ 0.1 are colored. NES, Normalized Enrichment Score. **F)** GSEA summary for HALLMARK, REACTOME, KEGG, and BIOCARTA gene sets in Combo treatment versus REST (Veh/DAC/TAZ) after 72 hours of treatment. Gene sets that meet a significance threshold of FDR ≤ 0.1 are colored. **G)** GSEA summary of upregulated BIOCARTA(B) and KEGG(K) gene sets that relate to calcium signaling (calcium transporters (dilated cardiomyopathy); calcium signaling associated pathways (TCR, Calcineurin, NFAT)). Gene Ratio is the ratio of the core enrichment genes for Combo versus REST for the indicated gene set divided by the total number of genes present in the gene set. **H)** GSEA analysis of Combo versus REST (Veh/DAC/TAZ) for Transcription Factor Targets (TFT) genes sets. **I)** Row z- score heatmaps of significantly upregulated transcription factors and T-cell receptor associated genes following combination inhibitor treatment of HCT116 for 72 hours. **J)** MA plot for differentially expressed genes in Combo treated HCT116 following 72 hours of treatment. Core enrichment genes identified from GSEA analysis of the indicated gene sets are colored.

Next, we determined which pathways were altered following the prolonged treatment using Gene Set Enrichment Analysis (GSEA). Consistent with previous studies of DNMT inhibition in colon, breast, and ovarian cancer, combination epigenetic therapy significantly upregulated pathways involved in extracellular matrix (ECM) organization (37,97) and a viral mimicry-associated innate immune response (6,8,37) (Figure 5E, **Figure S5F**). Although upregulation was the predominate direction of the transcriptional response, GSEA revealed that MYC and its targets were consistently downregulated with combination treatment (Figure 5E, **Figure S5G-H**), a response previously reported with combination DNMT and histone deacetylase (HDAC) inhibitor treatment (66,98). Taken together, EZH2 inhibition strongly enhanced the appreciated transcriptional re-programming effects of DNMT inhibition following prolonged drug exposure, including therapeutically actionable effects of innate immune pathway activation and MYC pathway downregulation.

### Acute transcriptional responses to combined DNMT and EZH2 inhibition reveal upregulation of calcium-calcineurin-NFAT pathway signatures and PRC2 target gene expression

As was observed at the prolonged treatment time point, GSEA of transcriptional signatures unique to acute combination treatment revealed induction of innate immune response pathways, extracellular matrix organization, and known PRC2 target genes (Figure 5F, **Figure S5I**). Notably, among the most significantly upregulated pathways following acute combination treatment were gene sets associated with T-cell activation and calcium signaling (Figure 5F-G). Converging on this observation, and from motif analysis described in Figure 4I-L, AP-1 (composed of JUN/FOS), MEF2, and NFAT target genes were among the most enriched gene sets upregulated with acute combination treatment (Figure 5H), and expression of these transcription factors was stimulated by the drug combination (Figure 5I). Moreover, many NFAT target genes, genes associated with T-cell receptor signaling, and genes involved in calcium signaling were induced from a silenced state following acute combination treatment (Figure 5J). Transcriptional upregulation of IFI27 (Interferon alpha-inducible protein-27, a downstream component of the viral mimicry pathway and a combo-responsive gene) and components of the calcium-calcineurin-NFAT signaling pathway in response to the drug combination was validated in several CRC cell lines (**Figure S5I**). Collectively, acute combination treatment stimulated many of the transcriptional responses (i.e. TE expression, innate immune pathway activation) that become predominant after prolonged treatment but also revealed that calcium signaling, downstream calcium-associated transcription factors, and their targets were significantly upregulated, suggesting that changes to the chromatin environment mediated by the drug combination facilitate the activation of this pathway.

### Reduced DNA methylation and H3K27me3 at promoters is associated with acute transcriptional responses to the epigenetic drug combination

We next sought to define the relationship between chromatin modification changes and transcriptional responses observed with the combination drug treatment. To do this, we integrated siQ-ChIP (H3K27me3), EM-seq (DNA methylation), and available ENCODE data for EZH2, H3K4me3 and H3K27ac occupancy across promoters of genes that were significantly upregulated following acute drug exposure (Figure 5D, **S5D**). Genes upregulated in response to DAC (Figure 5D, all genes in red circle) had low levels of H3K27me3, H3K4me3, and H3K27ac, and high levels of DNA methylation across their TSS’s in vehicle-treated HCT116 cells (Figure 6A-B and **Figure S6A**). H3K27me3 levels were marginally elevated in this gene set following DAC treatment, consistent with a lack of EZH2 association (Figure 6A, **S6A**). Genes upregulated in response to TAZ (Figure 5D, all genes in green circle) had high levels of H3K27me3 across their promoters and DNA hypomethylation at their TSS’s (Figure 6A-B and **Figure S6A**). Notably, TAZ responsive genes appear to be in a “poised” bivalent state with both EZH2 and H3K4me3 flanking the TSS. Unlike DAC responsive genes, TAZ responsive genes showed elevated H3K27me3 following DAC treatment, suggesting that while these genes are primarily silenced by H3K27me3, DNA hypomethylation stimulates poised EZH2 to deposit additional H3K27me3 to maintain a repressed state.

**Figure 6.**
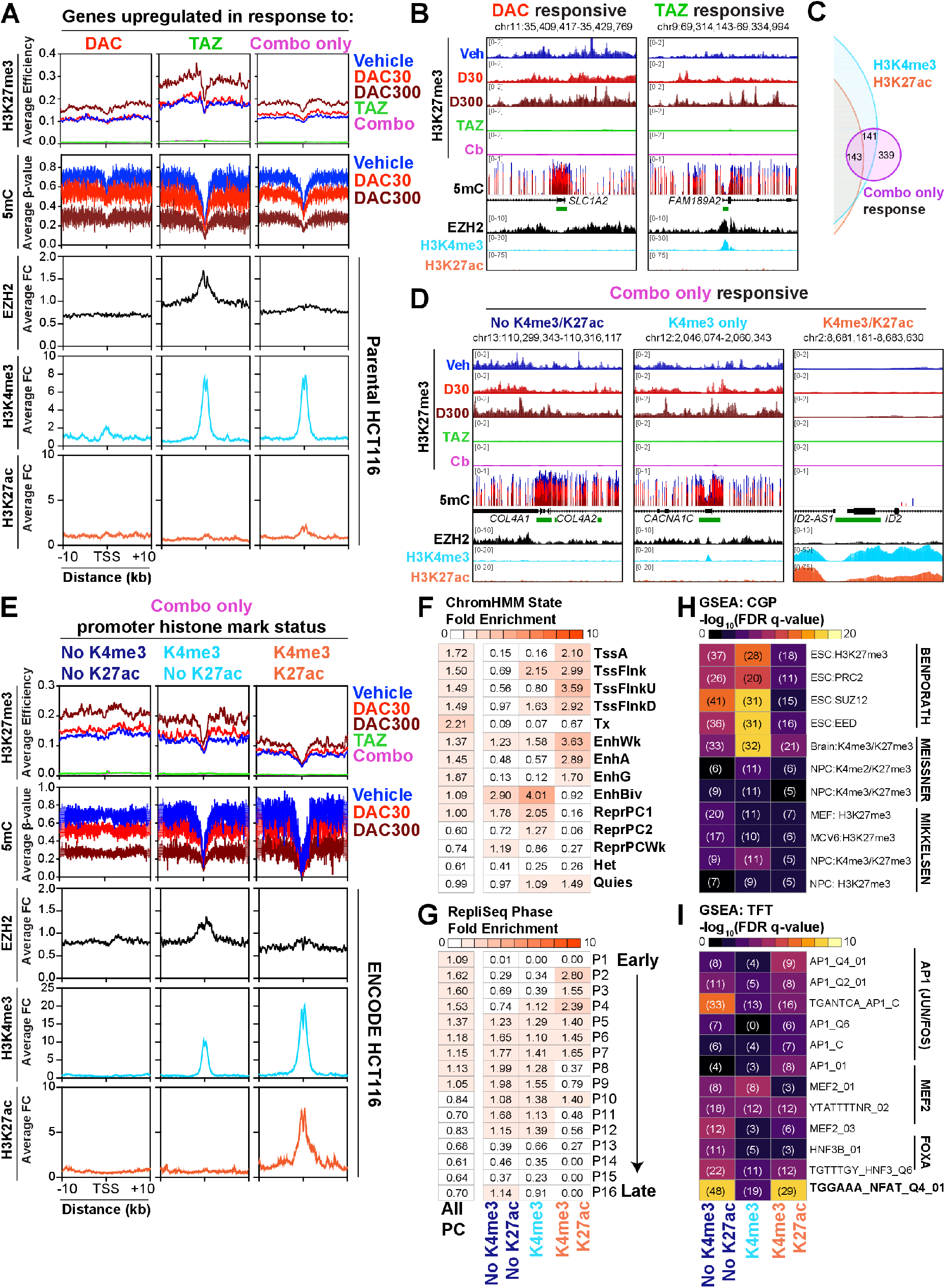
DAC plus TAZ combination treatment promotes acute transcriptional responses by inducing loss of DNA methylation and H3K27me3 at promoters. **A)** Integrative epigenomic analysis (rows) centered on the TSS of genes upregulated by the indicated inhibitor treatments (columns) in HCT116 following 72-hour treatment with 30 nM DAC and 1 µM TAZ. Average profiles for H3K27me3 and DNA methylation were determined from biological duplicate siQ-ChIP-seq and EM-seq measurements, respectively. EZH2, H3K4me3, and H3K27ac profiles were determined from publicly available ENCODE data for HCT116. **B)** Representative browser shots for a DAC-responsive locus (*SLC1A2)* and a TAZ-responsive locus (*FAM189A2*) demonstrating patterns observed in average profiles from **A. C)** Venn diagram of genes upregulated by combination treatment overlapped with genes that contain activating histone PTMs (H3K4me3/H3K27ac) in the promoter of their TSS (+/- 1000 bp). Combo only responsive genes are subdivided by the presence/absence of active histone PTMs in their promoters. **D)** Representative browser shots for the different subdivisions of promoter architecture of Combo only responsive genes derived from **A** and **C**. **E)** Integrative epigenomic analysis (rows) centered on the TSS’s of genes upregulated by the combination inhibitor treatment only (Combo only) subdivided by promoter histone PTM status from **C**. Average profiles for H3K27me3 and DNA methylation were determined from biological duplicate siQ-ChIP-seq and EM-seq measurements, respectively. EZH2, H3K4me3, and H3K27ac profiles were determined from ENCODE data publicly available for HCT116. **F-G)** Fold enrichment overlap analysis of genes upregulated by Combo only subdivided by the promoter histone PTM status with **D)** ChromHMM states and **E)** replication timing (RepliSeq) phases in HCT116. **H-I)** Hypergeometric overlap analysis of Combo only upregulated genes subdivided by promoter histone PTM status with **H)** GSEA C2: Chemical and Genetic perturbations related to known studies of PRC2 regulation and **F)** GSEA C3: TFT transcription factor targets. Heatmaps represent the significance of the overlap, number of genes in overlap provided in parentheses.

We next focused on the epigenetic architecture surrounding promoters of genes that exclusively responded to the combination treatment (Figure 5D, n=623 genes in purple circle). Unlike single agent responsive genes, combination only responsive genes did not demonstrate a clear overarching epigenetic regulatory mechanism (**Figure S6A**). Rather, genes exclusively upregulated by the combination treatment could be further subdivided into three groups based on promoter architecture of histone PTMs associated with active transcription (Figure 6C-D, **S6B**). Approximately half these genes were not marked by H3K27ac or H3K4me3. The remainder were marked by H3K4me3 alone, or in combination with H3K27ac (Figure 6C-D, **S6B**).

The expression of DAC- and TAZ-responsive genes is associated with drug-induced losses of DNA methylation and H3K27me3, respectively. This was also observed within the combination treatment responsive gene cluster lacking H3K4me3 and H3K27ac, where loss of both DNA methylation and H3K27me3 correlated with gene expression (Figure 6E). These genes were primarily found in Polycomb regulated regions of the genome (EnhBiv, ReprPC1, ReprPCWk) and in the transition regions from early to late replication timing (Figure 6F-G). Significantly, these genes were enriched not only for known Polycomb target genes in embryonic stem cells (Figure 6H) but also for genes that contain AP-1 (Jun/Fos) and NFAT transcriptional factor binding motifs (Figure 6I).

Similar to the TAZ responsive gene cluster, the combination treatment cluster marked by H3K4me3 had bivalent promoter architecture with H3K27me3 and EZH2 across the TSS (Figure 6E, middle column). These bivalent genes were also found in regions of the genome known to be associated with Polycomb regulation (EnhBiv, ReprPC1, ReprPC2) and the transition regions from early to late replication timing (Figure 6F,G). Consistent with a bivalent promoter architecture, this subgroup of genes was significantly enriched for known Polycomb target genes in embryonic stem cells (Figure 6H) but did not demonstrate strong enrichment for transcription factor binding sites in the promoter (Figure 6I).

Finally, the combination treatment cluster marked by H3K4me3 and H3K27ac showed no significant changes in H3K27me3 (lowest signal among all combination only responsive genes) and the deepest depletion of DNA methylation at the TSS among all treatments, epigenetic signatures consistent with active gene transcription (Figure 6E, right panel). Indeed, genes in this subcategory were already active in vehicle treated HCT116 cells, and they were further upregulated by the combination treatment (**Figure S6C**), suggesting that upregulation of these genes is a downstream effect of activation of other upstream genes and not dependent on changes in local epigenetic architecture induced by the drug combination. Consistent with this notion, these genes were not located in known Polycomb regulated regions and instead were enriched for known active regions of the genome and earlier replication timing phases (Figure 6F,G). Additionally, these genes were not enriched for known Polycomb targets (Figure 6H); however, they did demonstrate enrichment for targets of AP-1 (Jun/Fos) and NFAT transcription factor binding motifs (Figure 6I), suggesting that upregulation of these genes is a downstream effect of calcium signaling and associated transcription factor activation by the combination treatment.

### Calcium-calcineurin-NFAT pathway inhibition limits the transcriptional activating effects of combined DNMT and EZH2 inhibition

Building from the observation that the acute DNMT and EZH2 inhibitor combination stimulates expression of gene sets related to the Calcium-calcineurin-NFAT signaling pathway, and that genomic loci that undergo an “epigenetic switch” from DNA methylation to H3K27me3 when treated with DAC are enriched for AP-1 and NFAT binding motifs, we sought to define the relationship between calcium signaling and transcriptional regulatory effects of this epigenetic drug combination. Previous work in CRC lines showed that silencing of *SFRP1* and other TSGs with promoter CGI hypermethylation can be reversed by stimulating calcium signaling with cardiac glycosides (99). Consistent with this observation, the synergy between DNMT inhibition and EZH2 inhibition in stimulating *SFRP1* expression was blocked in the presence of calcium signaling antagonists, including a lymphocyte-specific protein tyrosine kinase (LCK) inhibitor (blocks calcium signaling at the plasma membrane), the calcium chelator EGTA, and the calcineurin inhibitor Cyclosporin A (CsA) (Figure 7A).

**Figure 7:**
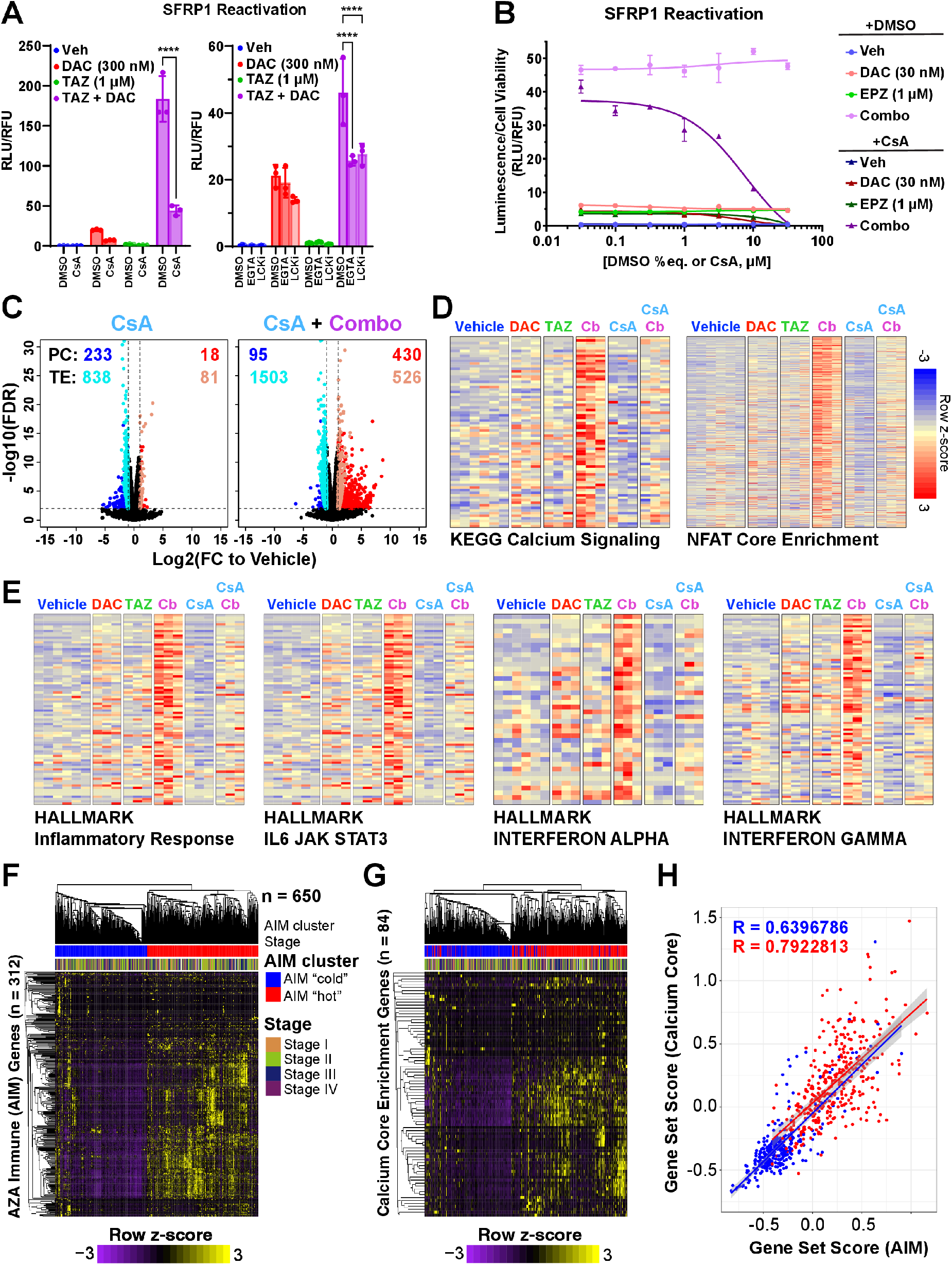
The transcriptional activating effects of the DAC plus TAZ combination treatment requires signaling through the calcium-calcineurin-NFAT pathway. **A-B)** *SFRP1* luciferase reporter activity measurements following 72-hour treatments with DAC and/or TAZ in the presence of calcium signaling antagonists. Data are presented as mean +/- SD of technical triplicates. **C)** Volcano plots for PC genes and TEs in HCT116 following 72-hour treatment (TAZ: 1 µM, DAC: 30 nM, CsA: 5 µM). Data points are plotted as averages from three biological replicates. Transcripts that meet a significance threshold of Log_2_(FC) ≥ 1.0 and FDR ≤ 0.01 are colored. Black points are PC transcripts that do not meet threshold conditions. **D)** Row z-score heatmaps of core enrichment genes from GSEA analysis of Combo versus REST (Veh/DAC/TAZ) for the indicated gene sets (left: KEGG Calcium Signaling; right: TFT NFAT targets). **E)** Row z-score heatmaps of core enrichment genes from GSEA analysis of Combo versus REST (Veh/DAC/TAZ) for the indicated gene sets. **F)** Supervised heatmap clustering of colon AIM gene expression among TCGA COAD READ tumor patients (n = 650 patients). AIM “hot” cluster refers to patients that exhibit relative high expression of the colon AIM genes and AIM “cold” cluster refers to patients with relative low expression of the colon AIM genes. AIM cluster and Stage annotations are provided and carried through the subsequent analysis. **G)** Supervised heatmap clustering of KEGG Calcium Signaling core enrichment (derived from 72-hour Combo vs REST GSEA analysis) gene expression among TCGA COAD READ tumor patients. Annotations are carried from **F. H)** Pearson correlation analysis of AIM genes versus Calcium genes among TCGA COAD READ patients. Each dot represents a patient. Gene Set Scores (for AIM and Calcium) were calculated by averaging the z-scores across the indicated gene set for an individual patient.

CsA was most effective at blocking *SFRP1* expression and did so in a dose-dependent manner (Figure 7A-B). In response to intracellular calcium influx, calcineurin, a calcium/calmodulin-dependent serine/threonine protein phosphatase, promotes NFAT translocation to the nucleus by dephosphorylating a nuclear localization signal (100–102). CsA inhibits calcineurin, and thus NFAT translocation, by forming a complex with cyclophilins that bind calcineurin and prevent its phosphatase activity. Although reported to induce G0/G1 cell cycle arrest (103), a concentration of CsA (5 µM) that robustly blocked both *SFRP1* expression (Figure 7B) and NFAT accumulation in the nucleus (**Figure S7A**) had no noticeable effects on the fraction of HCT116 in each cell cycle phase (**Figure S7B**) nor did it prevent the activity of the epigenetic drugs (**Figure S7C,D**). Collectively, these data suggested that calcium signaling pathway induction, an acute transcriptional response to the DNMT inhibitor and EZH2 inhibitor combination, contributes to downstream transcriptional responses to these epigenetic drugs.

### Calcium-calcineurin-NFAT signaling is required for the viral mimicry response induced by combined DNMT and EZH2 inhibition

We next performed RNA-seq on HCT116 acutely exposed to the DNMT and EZH2 inhibitor combination in the presence of CsA. CsA alone downregulated the expression of a limited number of PC genes and TEs (233 and 838, respectively) while also preventing the upregulation of many combo- responsive genes (Figure 7C). CsA reduced the transcriptional activating effects of the epigenetic drug combination to 430 PC and 526 TE transcripts (Figure 5C, right panel and Figure 7C, right panel). Consistent with calcineurin being a major target of CsA, genes involved in calcium signaling were not induced by the epigenetic drug combination when CsA was co-administered (Figure 7D, left panel). In addition, CsA blocked the expression of NFAT target genes induced by the DNMT and EZH2 inhibitor combination (Figure 7D right panel).

The effect of CsA on TE induction by the epigenetic drug combination prompted us to consider whether calcium signaling pathway induction was needed for stimulating the expression of genes associated with an innate immune response to viral infection. Indeed, the expression of genes associated with inflammation, IL6/JAK/STAT, interferon alpha, and interferon gamma signaling were attenuated when calcium pathway signaling was blocked with CsA (Figure 7E).

Finally, we asked if this strong link between intracellular calcium signaling activation and the innate immune response observed in our cancer cell line models could be observed in biopsied human tumors. To conduct this analysis, we leveraged a set of known innate immune response genes that consistently respond to low-dose AZA treatment across a panel of colorectal cancer cell lines (104) and asked if this gene set could cluster TCGA COAD READ patients (Cancer Genome Atlas Network, 2012a) based on innate immune response. Indeed, using this <u>A</u>ZA <u>IM</u>mune gene set (AIM), we were able to reproduce the separation of COAD READ patients into AIM ‘hot’ (high innate immune response) and AIM ‘cold’ (low innate immune response) clusters (Figure 7F). Next, we annotated the COAD READ patients by AIM ‘hot’ or ‘cold’ response (Figure 7F) and then, using our core enrichment KEGG Calcium Signaling gene set (Figure 7D), we determined that AIM ‘hot’ COAD READ patients also had high expression of calcium signaling genes (Figure 7G). Indeed, Pearson correlation analysis of the average calcium gene response to the average innate immune response among individual patients measured a significant association between these two expression profiles (Figure 7H). In addition to colon cancer-specific AIM genes, we also profiled breast and ovarian cancer specific AIM gene sets (104) among TCGA BRCA (**Figure S7E-G**) (106) and TCGA OV (**Figure S7H-J**) (107) patients, respectively. Consistent with colorectal cancer patients, breast and ovarian cancer patients also had a significant correlation between expression of calcium signaling genes and the innate immune response (**Figure S7G, J**), further emphasizing the connection between these two pathways.

## Discussion

### Therapeutic implications of combined DNMT and EZH2 inhibitor therapy

In this study, we show that compensatory accumulation of H3K27me3 after DAC treatment occurs primarily at distal Polycomb regulated regions, preventing full transcriptional activation by DNMT inhibitors. This H3K27me3 accumulation therefore represents a therapeutic vulnerability in colorectal cell lines that are normally insensitive to single agent EZH2 inhibition. The unmet potential of DNMT inhibition therapy in patients with solid tumors underscores the importance of our findings.

Previous studies have attempted to exploit the crosstalk between DNA methylation and histone PTMs to treat cancer (108,109). For example, DNMT inhibitors have been combined with HDAC inhibitors as a way to increase chromatin accessibility and activate otherwise silenced TSGs (13,70,108,110,111). Combined DNMT and HDAC inhibitor treatments have had mixed effects on clinical response rates and survival benefit compared to single agent DNMT inhibitor treatment (112–118). HDAC inhibition was also associated with increased DNA damage and this could explain some resultant pharmacodynamic antagonism with DNMT inhibition in cancer cell lines and patients (25,116,117,119–124).

Our study suggests simultaneously targeting repressive lysine methylation signaling at H3K27, particularly using the clinically applied EZH2 inhibitor TAZ, represents a different approach to augment the transcriptional and therapeutic effects of DNMT inhibition. Recent studies have begun examining the combinatorial use of DNMT and EZH2 inhibitors, with reports of cooperative therapeutic benefit that include reduced xenografted tumor or cell line growth, antineoplastic action, increased T-cell infiltration, re-sensitization to immunomodulatory drugs, re-expression of silenced TSGs, and activation of the viral mimicry response (11,54,56–59,109). Consistent with these findings, perturbing DNA methylation, genetically or as a consequence of chemotherapy/metabolic perturbation, renders cancer cells sensitive to EZH2 inhibition. Conversely, tumors with genetically compromised PRC2 activity are hypersensitive to DNMT inhibition (53,55,125,126). Our studies build from these reports, corroborating the combination anti-proliferative effects of DNMT and EZH2 inhibition in colorectal cancer models while also providing in-depth analysis of the molecular effects of combined DNMT and EZH2 inhibition on the epigenome and transcriptome, the importance of which is discussed below. We also inform on the rational selection of epigenetic inhibitors used in clinical practice and provide rationale for expanding the use of EZH2 inhibitors beyond tumor types that show a dependency on PRC2.

### Immunomodulatory effects of combined DNMT and EZH2 inhibitor treatment

We and others found that TEs become activated after DNMT inhibition, which triggers a “viral mimicry response” mediated through dsRNA sensing and interferon signaling (6,8). Here, we show that TAZ enhances the DAC-induced expression of TEs and innate immune response pathways associated with viral mimicry. Activating antiviral response programs in tumors is associated with a better response to immunotherapy in preclinical models (10–12), so these data suggest future directions for testing whether DAC + TAZ treatment might improve tumor response to immunomodulating agents.

Combined DAC and TAZ treatment also transcriptionally upregulated components of the calcium-calcineurin-NFAT signaling pathway in numerous colon cancer cell lines (Figure 5F-J and **S5I**), but it is still unclear how this influences oncogenesis or solid tumor response to treatment (99,127,128). The calcium-calcineurin-NFAT pathway is instead primarily associated with T-cell activation (101,129), and it is known that single agent DNMT or EZH2 inhibitor therapy activates T-cells and can improve immune modulation in cancer cells and *in vivo* cancer therapy models (10,12,130–134). Our data are consistent with a model whereby combined DNMT and EZH2 inhibitor treatment stimulates the tumor cell-extrinsic immune microenvironment, either by inducing viral mimicry or by activating the calcium-calcineurin-NFAT pathway. This may be of particular relevance to microsatellite stable (MSS) colon tumors that respond poorly to immune checkpoint inhibitors (135). The MSS cell lines Colo205 and CACO2 showed both antineoplastic effects and transcriptional enrichment of calcium signaling following DAC and TAZ combination treatment (Figure 3B and **S5I**). Future studies will consider whether the DAC and TAZ combination helps overcome immunotherapy resistance in MSS colon cancers.

### Rational EZH2 inhibitor selection

We were surprised to find that the two most potent EZH2 inhibitors (TAZ and CPI-1205) were the least toxic of all EZH2 inhibitors tested in this study. The disconnect between cell viability and potency for these toxic compounds suggests these molecules have off-target activities. Indeed, recent studies have called for re-evaluating the off-target effects of small molecules used experimentally and in clinical trials (including the EZH2 inhibitor UNC1999 used in this study), suggesting the mechanism of action behind their efficacy and/or toxicity may stem from different interactions than predicted (136,137).

Importantly, compounds like TAZ and CPI-1205 showed characteristics of low toxicity coupled with high potency also synergized the best with DAC. The limited single agent toxicity of select EZH2 inhibitors may be key for maximizing combination effects with DNMT inhibitors that require DNA synthesis for drug action. Given the broad interest in EZH2 inhibition in both laboratory and clinical settings (67–69), this study provides valuable insight as to how to rationally evaluate and assay these small molecules, including factors outside of cytotoxicity.

### Relationship between epigenetic and transcriptional responses

The concept of an “epigenetic switch” in which the loss of one repressive mark induces compensation by a different silencing mechanism is not new to this field (46,47,138,139). In this study, we determined that an “epigenetic switch” between DNA methylation and H3K27me3 does not occur on a global scale, but rather at specific Polycomb regulatory regions of the genome such as bivalent enhancers. Our study revealed a common theme for where the “epigenetic switch” occurs in which loss of DNA methylation at genomic regions with low basal levels of H3K27me3 induced increases in the repressive mark due to pre-existing EZH2 occupancy. Notably, this reversion to PRC2-mediated repression following the removal of DNA methylation has been observed in normal developmental contexts where H3K27me3 helps maintain transposon repression after genome-wide induction of DNA methylation loss in mouse embryonic stem cells (140), adding evidence for the targetability of the interconversion of these two distinct silencing marks. Our combination treatment effectively blocked the “epigenetic switch” that occurs with single-agent DAC treatment, leading to accessibility of calcium signaling-associated transcription factor binding motifs (AP-1, NFAT) for downstream regulation of transcriptional responses.

Early genome-wide studies reported anti-correlations between DNA methylation and H3K27me3 distributions leading to the paradigm that these marks are mutually exclusive (46,139). However, we and others report that these marks can indeed co-exist and silence gene transcription in a cooperative manner (58,95,96). In this study, we identified over 300 PC genes (including TSGs such as *SFRP1*) that were dually silenced by both DNA methylation and H3K27me3 at promoter regions in HCT116 cells. Importantly, single agent treatment with low-dose DAC or TAZ did not reactivate expression of these genes. Rather, the combination of DNMT and EZH2 inhibition was required to effectively remove both DNA methylation and H3K27me3, respectively, to permit expression of these genes. Our results are consistent with the observation that cancer cells are prone to co-existence of these modifications and that combination DNMT and EZH2 inhibitor treatment can effectively target genes that are dually modified for activation.

## Acknowledgements

We thank members of the Rothbart and Baylin laboratories, Peter Laird, Ronen Marmorstein, and members of the Epigenetic Therapies SPORE for helpful discussions. We also thank Kelly Foy and Nicole Vander Schaaf for assisting with harvesting intestinal adenomas from APC^Min+^ mice, Ben Johnson and Josh Jang for discussions on bioinformatic analyses, and Darrell Chandler for critical reading of the manuscript. We acknowledge support from the Van Andel Institute Genomics Core, Bioinformatics and Biostatistics Core, Flow Cytometry Core, and Optical Imaging Core. This work was supported by grants from the National Institutes of Health to S.B.R (R35GM124736, P50CA254897), S.B.B. (P50CA254897), A.A.C. (F32CA225043), and grants from the American Cancer Society to S.B.R. (RSG-21-031-01-DMC) and ACS Michigan Cancer Research Fund to R.L.T. (PF-16-245-01- DMC).

---

**Figure S1 (Related to Figure 1). DAC synergizes with TAZ better than other EZH2 inhibitors to induce expression of an epigenetically silenced TSG A)** Western blot analysis of H3K27me3 from HCT116 following 72-hour exposure to TAZ or siRNA-mediated knockdown of PRC2 components with or without DAC. Representative short (top) and long (middle) exposures for H3K27me3 are shown. **B)** Chemical structures of SAM-competitive EZH2 inhibitors used in this study. **C-D)** NLuc reporter activity measurements following 72-hour treatment with TAZ, **(C)** its precursor molecule EPZ005687, or **(D)** its derivative EPZ011989 with or without a fixed concentration of DAC (30 nM). Error bars are presented as mean +/- SD of technical triplicates. **E)** Density plots of β-value distributions for all CpG probes (n = 705,169) that passed quality control analysis on Illumina MethylationEPIC arrays from the indicated drug treatments (biological duplicate) in HCT116. β-values range from [0-1] with 0 indicating no DNA methylation and 1 indicating complete DNA methylation. **F)** Dose-response High Resolution Melt (HRM) analysis of DNA methylation at the indicated loci following 72-hour treatment of HCT116. RPL30, unmethylated control gene. SFRP1, methylated CpG island promoter. Data are presented as mean +/- SD of technical duplicates. **G)** Surface plot and Bliss synergy score using SynergyFinder2.0 to analyze combination DAC plus TAZ dose response curves derived from the *SFRP1*-NLuc reporter assay after 6 days of treatment following a single application of drugs. **H)** qRT-PCR validation of the *SFRP1* transcriptional response in (left panel) HCT116 DNMT1-hypomorph (MT1) cells or (right panel) wild-type HCT116 cells treated for 72 hours with the indicated drugs. 2^-ΔΔC(t)^ normalized to housekeeping gene RPL4 and no treatment or vehicle. Data are presented as mean +/- SEM of technical duplicates (MT1) or +/- SD biological triplicates (WT). ****p<0.0001, ***p<0.0005, **p<0.004.

**Figure S2 (Related to Figure 2). Single agent EZH2 inhibitor potency and therapeutic efficacy do not correlate in HCT116 A)** Western blot analysis of H3K27me3 from HCT116 cells following 72-hour exposure to the indicated concentrations of EZH2 inhibitors. Concentration ranges of EZH2 inhibitors vary by inhibitor (indicated by corresponding wedge sizes) to obtain comparable levels of H3K27me3 loss. **B)** H3K27me3 levels from **(A)** were quantified by relative band intensity using ImageJ densitometry and normalized to both the relative band intensity of the H3 loading control and as a percent of vehicle (DMSO). **C)** *In vitro* PRC2 activity assays with recombinant mononucleosome substrates in the presence of the indicated EZH2 inhibitors. Data are presented as mean +/-SEM of technical triplicates. **D)** Dose response viability curves (CellTiter-Glo normalized to vehicle) for wild-type HCT116 cells exposed to the indicated EZH2 inhibitors for 72 hours. Data are presented as mean +/- SD of technical triplicates.

**Figure S3 (Related to Figure 3). Concentration-dependent therapeutic effects of combined DNMT inhibitor plus EZH2 inhibitor treatment across colon cancer cell lines A)** Western blot analysis of H3K27me3 following 72-hour TAZ exposure across a panel of colon cancer cell lines. NTx, no treatment. Representative short (top) and long (middle) exposures for H3K27me3 are shown. **B)** CellTiter-Fluor viability assays for Colo205 and HCT15 cell lines after 72-hour treatment with the indicated drugs. Data are presented as mean +/-SD of technical triplicates. **C)** High Resolution Melt analysis of the indicated colon cancer cell lines treated with Vehicle or DAC at indicated concentrations for 72 hours. *RPL30*, unmethylated control gene. *SFRP1*, methylated CpG island promoter. Data are presented as mean +/- SD of technical duplicates. **D)** Confluency measurements from Incucyte imaging of the indicated cell lines. Cells were split and drugs were refreshed every 72 hours for the indicated number of days prior to plating (“pretreatment”) and were re-treated and re-plated again at Day 0 of Incucyte measurements. Data are presented as mean +/- SEM of technical well triplicates averaged from 9 images per timepoint and treatment. **E-G)** Confluency measurements from Incucyte imaging of HCT116 treated with the indicated drugs at Day 0. Data are presented as mean +/- SEM of technical well triplicates averaged from 12 images per timepoint and treatment. **H)** Representative images of tumoroids counted in Figure 3D. Type 1: healthy, spherical with defined lumen; Type 2: spherical, clustered with visible lumen; Type 3: unhealthy, clustered with dark lumen; and Type 4: small, singular with no discernible lumen.

**Figure S4 (related to Figure 4). EPIC array, ChIP-EM-seq quality control, and custom ChromHMM build A)** Density scatterplots of Illumina MethylationEPIC CpG probe (n = 704,169) β-values comparing different treatments. Pearson correlations (R) for CpG probe β-values between drug treatment samples were calculated. **B)** Histogram of Δβ-values (change in DNA methylation) relative to Vehicle samples of highly methylated CpG probes (Average Vehicle β-value ≥ 0.85, n = 442,383). β- values were averaged across biological duplicates for each drug treatment and then the Δβ-value (relative to Vehicle) was calculated. **C)** Upset genomic annotation plot of shared hypomethylated Differentially Methylated Regions (DMRs) between DAC and DAC plus TAZ combination treatment (n = 8,998). DMRs are defined as regions with at least 5 contiguous CpGs (by the EPIC array) with a mean Δβ-value of 0.15. The top 4 predominant genomic annotations are provided. **D)** PCA analysis of siQ-ChIP libraries (left panel) and EM-seq libraries (right panel). All experiments were conducted in biological duplicate. **E)** Quality control analysis for enrichment of H3K27me3 fragments and subsequent read coverage by EM-seq. All PC genes were clustered based on k-means (n=4) analysis of H3K27me3 efficiency across all samples at and surrounding the TSS (top panel). The average read coverage from EM-seq was then calculated for all H3K27me3 clusters. **F)** Percent cytosine methylation of CpG and CpH (H = A,T,or C) dinucleotides in EM-seq data. **G)** Violin box plot of H3K27me3 efficiency z-score categories versus average DNA methylation (100 bp bins). All 100 bp bins were stratified by H3K27me3 efficiency from highest to lowest, and then the z-score was calculated for each bin. H3K27me3 efficiency z-score categories are as follows: lowest (z-score = [-1,0]), n = 1,821,894 100 bp bins; low (z-score = [0,1]) n = 1,261,814 100 bp bins; high (z-score = [1-2]), n = 508,308 100 bp bins; highest (z-score ≥ 2), n = 428,846 100 bp bins. **H)** Custom ChromHMM emission state profile for HCT116 built from H3 Histone PTMs (ENCODE datasets: H3K4me1, H3K27ac, H3K4me3, H3K36me3; Custom datasets: H3K9me3, H3K27me3). Emission state names were assigned based on the predominant PTM(s) present. **I)** Genome coverage (bp) of each custom ChromHMM emission state for HCT116 from panel **H**. **J)** Enrichment analysis comparing distributions of custom ChromHMM emission states with 16-phase Repli-seq (replication timing) regions. Note that Polycomb regulated regions (EnhBiv, ReprPC1, ReprPC2) are primarily enriched in intermediate replication timing phases and ReprPCWk is enriched in late replication timing phases. **K-L)** Average profiles and heatmaps for (**K**) H3K27me3 efficiency and (**L**) EZH2 occupancy at conserved genomic regions that show increased H3K27me3 following 30 nM DAC treatment (n=3,223 peaks).

**Figure S5 (related to Figure 5). RNA-seq QC and GSEA analysis A)** Principal Components Analysis (PCA) of total RNA-seq conducted on HCT116 treated with Vehicle (DMSO), 30 nM DAC, 1 µM TAZ, or the combination for 6 days (2 doses for all treatments). All treatments were conducted in biological triplicate. Proportion of variance in each principal component is displayed below. **B)** Row z-score heatmaps of significantly upregulated genes (Log_2_(FC) ≥ 1.0 and FDR ≤ 0.01) for single agent DAC treatment (DAC Response), TAZ treatment (TAZ Response), and combination treatment (Combo only response) (see Figure 5B). **C)** Principal Components Analysis (PCA) of total RNA-seq conducted on HCT116 treated with Vehicle (DMSO), 30 nM DAC, 1 µM TAZ, or combination treatment for 72 hours. All treatments were conducted in biological triplicate. Proportion of variance in each principal component is displayed below. **D)** Row z-score heatmaps of significantly upregulated genes (Log_2_(FC) ≥ 1.0 and FDR ≤ 0.01) for single agent DAC treatment (DAC Response), TAZ treatment (TAZ Response), and combination treatment (Combo only response) (see Figure 5D). **E)** Scatterplot of PC gene Log_2_(FC) values from Day 6 treatments versus Day 3 treatments in HCT116. **F)** GSEA analysis of HALLMARK gene sets associated with epigenetic therapy response 6 days post-treatment. Top: Row z-score heatmaps of core enrichment genes for the indicated HALLMARK gene set; Bottom: GSEA plots for Combo versus REST with the measured NES and FDR values reported. **G)** GSEA plots for HALLMARK MYC TARGETS V1/V2 for Combo versus REST with the measured NES and FDR values reported. **H)** GSEA analysis of Combo versus REST (Veh/DAC/TAZ) for Chemical and Genetic Perturbation genes sets associated with PRC2 target studies. **I)** Gene expression measured by qPCR from cell lines treated for 72 hours with the indicated drugs. 2^-ΔΔC(t)^ normalized to housekeeping gene RPL4 and no treatment or vehicle. Data are presented as mean +/- SD of biological duplicates or triplicates as indicated by individual dots. Genes are colored according to the subset they represent: NFAT targets (orange), Calcium signaling (grey), and Transcription Factors (light blue).

**Figure S6 (related to Figure 6). Integrative promoter-centric epigenomic analysis of the DAC plus TAZ combination A)** Integrative epigenomic analysis (columns) centered on the TSS of genes upregulated by the indicated inhibitor treatments (rows) in HCT116 following 72-hour treatment with 30 nM DAC and 1 µM TAZ. EZH2, H3K4me3, and H3K27ac profiles were determined from ENCODE data publicly available for HCT116. **B)** Integrative epigenomic analysis (columns) centered on the TSS of genes upregulated by the combination treatment alone (rows) in HCT116 cells following 72-hour treatment with 30 nM DAC and 1 µM TAZ. Upregulated genes were further subdivided based on presence or absence of activating histone marks (H3K27ac, H3K4me3) from Figure 6C. EZH2, H3K4me3, and H3K27ac profiles were derived from publicly available ENCODE data for HCT116. **C)** MA plot for differentially expressed genes in Combo only treated HCT116 colored by the associated promoter histone PTM architecture. Vehicle logCPM provides an indication of the level of expression of the gene in control cells with lowly expressed genes to the left and highly expressed genes to the right.

**Figure S7 (Related to Figure 7). Cyclosporin A disrupts the calcium-calcineurin-NFAT signaling axis in HCT116 but does not prevent epigenetic drug activity A)** Immunofluorescence staining of NFAT1 (green) and DAPI (red) in HCT116 treated with CsA for 72 hours and challenged with DMSO vehicle (left) or Ionomycin (right) for 60 min. **B)** HCT116 cell cycle analysis by proxy of propidium iodide staining and flow cytometry quantification after cyclosporin A (CsA) treatment for 72 hours. Data are presented as mean +/- SD of technical duplicates. **C)**. High Resolution Melt (HRM) analysis of HCT116 treated with Vehicle, DAC (30 or 300 nM), TAZ (1 µM) or Combo in presence or absence of CsA (5 µM) for 72 hours. RPL30, unmethylated control gene. SFRP1, methylated CpG island promoter. Data are presented as mean +/- SD of technical duplicates. **D)** Western blot analysis for H3K27me3 from HCT116 treated with TAZ and/or DAC in presence of CsA (5 µM) for 72 hours. **E)** Supervised heatmap clustering of breast AIM gene expression among TCGA BRCA tumor patients (n = 1111 patients). AIM “hot” cluster refers to patients that exhibit relative high expression of the breast AIM genes and AIM “cold” cluster refers to patients with relative low expression of the breast AIM genes. AIM cluster and Stage annotations are provided and carried through the subsequent analysis. **F)** Supervised heatmap clustering of KEGG Calcium Signaling core enrichment (derived from 72-hour Combo vs REST GSEA analysis) gene expression among TCGA BRCA tumor patients. Annotations are carried from **E. G)** Pearson correlation analysis of AIM genes versus Calcium genes among TCGA BRCA patients. Each dot represents a patient. Gene Set Scores (for AIM and Calcium) were calculated by averaging the z-scores across the indicated gene set for an individual patient. **H)** Supervised heatmap clustering of breast AIM gene expression among TCGA OV tumor patients (n = 421 patients). AIM “hot” cluster refers to patients that exhibit relative high expression of the ovarian AIM genes and AIM “cold” cluster refers to patients with relative low expression of the ovarian AIM genes. AIM cluster and Stage annotations are provided and carried through the subsequent analysis. **I)** Supervised heatmap clustering of KEGG Calcium Signaling core enrichment (derived from 72-hour Combo vs REST GSEA analysis) gene expression among TCGA OV tumor patients. Annotations are carried from **H**. **J)** Pearson correlation analysis of AIM genes versus Calcium genes among TCGA OV patients. Each dot represents a patient. Gene Set Scores (for AIM and Calcium) were calculated by averaging the z-scores across the indicated gene set for an individual patient.

## Methods

### Cell and tumoroid culture

HCT116, HCT15, COLO205, SW620, and CACO2 colon cancer cell lines were purchased from ATCC and maintained according to ATCC recommendations in McCoy’s (Gibco 16600-082), RPMI 1640 (Gibco 11875-093), RPMI 1640 (Gibco 11875-093), Leibovitz’s L-15 (ATCC 30-2008), and EMEM (ATC, 30-2003) Medium, respectively, supplemented with Fetal Bovine Serum (Millipore Sigma F0926, 20% for CACO2 and 10% for the other cell lines) and 1% penicillin/streptomycin (Life Technologies, 15140-122) at 5% CO_2_ and 37°C.

Tumoroids were derived from an APC^Min+^ mouse small intestine adenoma. After euthanasia with CO_2_, the small intestine was excised, opened longitudinally, and vigorously washed 3x in PBS. Aadenomas were excised, minced, suspended in PBS, and centrifuged at 300 x g for 5 min. Pellets were resuspended in Dispase (STEMCELL Technologies 07923) and incubated at 37°C for 10 min. Dissociated tissue was centrifuged again at 300 x g for 5 min, washed in DMEM (Life Technologies 11965092), passed through a 70 µm filter, and centrifuged again. Dissociated cells were washed once more with DMEM, centrifuged, and resuspended in a 50:50 mixture of Matrigel (Corning 356231):modified HITES Media [DMEM/F12 50/50 (Life Technologies 11330032) supplemented with 1x Glutamax (Gibco 35050061), 1% penicillin/streptomycin (Life Technologies 15140-122), insulin-transferrin-selenium mix (Gibco Invitrogen 41400-045), 10% Fetal Bovine Serum (Millipore Sigma F0926) with 1.6 ng/mL hydrocortisone (Sigma H0888) and 50 ng/mL murine EGF (Thermo Fisher PMG8041) added immediately before use]. 40 µl of this mixture was seeded into each well of a pre-warmed 48 well plate. This was incubated for 10 min at 37°C to polymerize the Matrigel and then covered with modified HITES media. Once tumoroids formed, they were split using cold media and gently trituration and then replated within 50:50 Matrigel:modified HITES covered by modified HITES media every 7-10 days.

### siRNA transfection

ON-TARGETplus siRNA SMARTpools (5 µl of 20 µM stock) targeting EZH1, EZH2, or SUZ12 (Dharmacon L-004217-00-0005, L-004218-00-0005, and L-006957-00-0010) and siRNA non-targeting control siRNA pools (Dharmacon D-001810-10-05) were mixed with Lipofectamine RNAiMax (5 µL, ThermoFisher 13778075) in a total of 200 µL Optimem (Life Technologies 31985-062) and incubated for 20 min. McCoy’s Media, with or without DAC and supplemented with 10% FBS but no antibiotics, was refreshed, and the siRNA/Lipid mix was added dropwise for a final siRNA concentration of 50 nM. Cells were collected 72 hours after transfection.

### Drug treatments

Tazemetostat (TAZ or EPZ6438, Selleck Chemicals S7128 and Cayman Chemical 16174), CPI- 1205 (Selleck Chemicals S8353) EPZ0011989 (Selleck Chemicals S7805), EPZ005687 (Cayman Chemical 13966), GSK126 (Selleck Chemicals S7061 and Cayman Chemical 15415), GSK343 (Selleck Chemicals S7164), GSK503 (Cayman Chemical 18531), UNC1999 (Caymen Chemical 14621), GSK- 3484862 (Chemietek CT-CSKMI-714), Cyclosporin A (CsA, Selleck Chemicals S2286), and Ionomycin (Cayman Chemical 11932) were dissolved in DMSO and stored at −20°C. 5-aza-2’-deoxycytidine (DAC, Sigma A3656) was dissolved in DMSO:PBS at a ratio of 1:150 and stored at −80°C. Each drug or % vehicle equivalent was applied to attached cells or tumoroids suspended in 50:50 matrigel:modified HITES media.

### Proliferation assays

To measure cell viability, cells were plated in 96-well plates at a density of 4000-5000 or 500- 700 cells/well (for a 3- or 7-day exposure to drugs, respectively), treated the following day, and assayed with either CellTiter-Glo (Promega G7570) or CellTiter-Fluor (Promega G6080) kits according to the manufacturer’s protocol. Measurements were taken using a BiotTek SynergyNeo Microplate Reader. Background signal from media-only wells was subtracted to obtain the final reported relative luminescence units (RLU, CellTiter-Glo) or fluorescence units (RFU, CellTiter-Fluor).

To measure outgrowth of colon cancer cells after prolonged treatments, cells were plated in a 6- well dish and drug was applied to adhered cells the following morning. After 72 hours, cells were collected and re-plated directly in fresh media and drug for another 72 hours, for a total of 3-6 days of “pre-treatment” depending on the cell line. Following pre-treatment, cells were plated into either 12-well (10,000-20,000 cells/well) or 96-well (500-700 cells/well) dishes and treated for a final time, which is represented as “Day 0” following pre-treatments on confluency graphs. From these samples, 9-12 images were captured for each well at each timepoint and % confluency calculated using a Sartorius Incucyte S3.

For prolonged treatment of tumoroids, media and drugs were refreshed every 3-4 days and tumoroids split in a 1:4 ratio (regardless of response to drug treatment) every 7-10 days. At 21 days, brightfield images from three wells per treatment were taken using the 4x objective of a Nikon Eclipse TS2R microscope. Tumoroids were categorized and counted based on morphology categories, and the counter was blinded to the tumoroid treatment.

### *SFRP1*-NanoLuciferase Reporter Assay

The CellTiter-Fluor assay was duplexed with the Nano-Glo Luciferase Assay System (Promega N1110) to measure NanoLuciferase activity in *SFRP1*-NLuc experiments. For these experiments, cells were plated in 96-well plates as above for the cell viability assays, and media and drugs were refreshed after 72 hours for the 6-day synergy measurements. To duplex the assays, CellTiter-Fluor was used at a 5x concentration preceding the Nano-Glo assay. NanoGlo reagent was applied according to the manufacturer’s protocol, and a SynergyNeo Microplate Reader was used to obtain relative luminescence units (RLU). After background subtraction from media-only wells, RLU were normalized to CellTiter-Fluor RFU to account for cell viability. This normalization was also used as the input for the synergy score calculations. The expected drug combination responses were calculated based on the Bliss independence model using SynergyFinder2.0 (81). Deviations between observed and expected responses with positive and negative values denote synergy and antagonism respectively.

### Immunofluorescence staining

HCT116 cells were grown on Lab-Tek chamber slides (Thermo Fisher 154534) and treated for 72 hours with CsA preceding a 1-hour Ionomycin challenge. Slides were rinsed in PBS, fixed with 4% PFA for 15 min, and rinsed again with PBS. PFA was neutralized and cells permeabilized with Permeabilization Buffer (PBS, 0.3% Tx-100, 200 mM glycine) for 20 min. Cells were incubated with blocking buffer (PBS, 2% BSA, 0.1% Tx-100, 10% goat serum) for 30 min followed by incubation with primary antibody against NFAT1 [1:100, CST5861, which was validated by western blot of HCT116 lysates with siRNA-mediated NFAT1 knockdown (not shown)] in antibody buffer (PBS, 0.1% BSA, 0.1% Tx-100) for 60 min. Cells were washed twice with antibody buffer, incubated with secondary antibody (1:200; Invitrogen, A-11034) for 30 min, and washed twice with antibody buffer. Slides were coverslipped with SlowFade Gold mounting medium (Thermo Fisher S36936) and imaged using the 40x objective of an Olympus BX51 microscope and the assistance of the VAI Optical Imaging Core.

### Flow cytometry

Cells treated with 5 µM CsA for 72 hours were collected by trypsinization and centrifuged at 300 x g for 10 min. Cell pellets were resuspended in 0.5 mL cold PBS, and 4.5 mL of cold 70% ethanol was added dropwise to cells while gently vortexing. Cells were fixed on ice for 2 hours, centrifuged at 500 x g for 10 mins, washed in PBS, and centrifuged again. Cell pellets were resuspended in 500 µL of PI staining solution (500 µL PBS, 100 µg/mL RNase A, 50 µg/mL propidium iodide) and incubated at 4°C for two hours. Flow cytometry and analysis was performed by the VAI Flow Cytometry Core using a CytoFLEX S Flow cytometer.

### Western blotting

Cells were lysed on ice in cold CSK lysis buffer [10 mM PIPES pH 7.0, 300 mM sucrose, 100 mM NaCl, 3 mM MgCl_2_, 0.1% Triton X-100, universal nuclease, and protease inhibitor cocktail (Roche cOmplete Mini tablets, EDTA-free)] for 30 min and then centrifuged 10 min at 10,000 x g to remove insoluble protein. Total protein in the supernatant was quantified by Bradford Assay (BioRad 5000006), denatured in SDS Loading Buffer, and boiled for 10 min. 2-5 µg of protein was size-separated by SDS-PAGE and transferred to a PVDF membrane. Membranes were blocked for one hour at room temp (PBS, 0.1% Tween-20, and 5% BSA), washed in PBST (PBS and 0.1% Tween-20), and incubated with H3 antibody (1:50,000; EpiCypher 13-0001) or H3K27me3 antibody (1:2,000; CST9733) in blocking buffer overnight at 4°C. The H3K27me3 antibody was chosen for its high specificity and selectivity among H3K27me3 antibodies profiled by our lab on histone peptide arrays (141). Membranes were washed in PBST and incubated in HRP-conjugated secondary antibody (1:10,000; Sigma, GENA934) for 1 hour at room temp. Membranes were washed again in PBST, incubated in ECL, and imaged with film. ImageJ densitometry was used to quantify band intensity.

### High Resolution Melt (HRM) Assay for DNA methylation

DNA was isolated from treated cell pellets using a DNeasy Blood & Tissue Kit (Qiagen 69504) according to manufacturer’s protocol. The EZ DNA Methylation Kit (Zymo D5002) was used to bisulfite convert 500 ng of DNA per sample according to the manufacturer’s protocol. Bisulfite converted DNA was eluted in 10 µL of M-elution buffer from the kit and brought up to 54 µL total with DNase-Free water. 5 µL of the bisulfite converted DNA was combined with 10 µL of Precision Melt Supermix for High Resolution Melt Analysis (BioRad 1725112), 2 µL of Forward and Reverse primers (2 µM stock) and brought to 20 µL with DNase-Free water. A BioRAD CFX Opus93 Real-Time PCR System was used to amplify the DNA at a 60°C annealing temp for 39 cycles and then perform a melt analysis from 65 to 95°C with 0.1°C/10 sec increments. The melt temp (Tm) at the maximum reported RFU value was reported for each amplicon. An amplicon from an unmethylated gene, *RPL30,* was used to control for bisulfite conversion. An amplicon in the hypermethylated CGI promoter of *SFRP1* was used to report on DNA methylation changes. *RPL30* primers: Forward, TAATTTAGAAGAGATAGAGAATAGGATAGGAATTTTAG and Rev, ACCATCTTAACGACTACTATTAATAAATAAACTCCTAC. *SFRP1* primers: Forward, AGGGGTATTTAGTTGTTGGTTTGTTG and Reverse, CTTCTACACCAAACCACCTCAATA.

### RNA isolation, cDNA synthesis, and qRT-PCR

TRIzol Reagent (Invitrogen 15596026) was added directly to adhered cells, collected, and stored at −80°C. After thawing, total RNA was extracted following the manufacturer’s protocol, and RNA was resuspended in DEPC Nuclease-Free water. The High-Capacity cDNA Reverse Transcription Kit (Applied Biosystems 4368814) with the addition of RNAse was used according to the manufacturer’s protocol to synthesize cDNA from 2 µg RNA. Using the KAPA SYBR FAST qPCR Kit (Roche 07959567001) according to the manufacturer’s protocol, qRT-PCR was performed on technical duplicates for each gene and sample and run on a BioRAD CFX Opus93 Real-Time PCR System. Data was analyzed using 2^-dd(Ct)^ method (142) where the fold change was determined by normalizing to the RPL4 housekeeping gene and a no treatment or vehicle treated sample.

### Chromatin Immunoprecipitation (ChIP)

Cells exposed to drugs for 72 hours were fixed in buffer (1% formaldehyde, 50 mM HEPES-KOH pH 7.6, 100 mM NaCl, 1 mM EDTA pH 8.0, 0.5 mM EGTA pH 8.0) for 10 min at room temp with shaking and then quenched with 125 mM glycine 5 min at room temp. Cells were scraped into cold PBS, washed 2X with cold PBS, flash frozen in liquid N2, and stored at −80°C until use. Thawed pellets were lysed in LB1 (50 mM HEPES-KOH pH 7.6, 140 mM NaCl, 1 mM EDTA, 10% Glycerol, 0.5% NP- 40, 0.25% Triton X-100, Roche protease inhibitor cocktail) for 20 min with rotation at 4°C and cleared by centrifugation at 300 x g for 5 min at 4°C. Supernatant with intact nuclei was set aside. Cell pellets were lysed again in 4x LB1 (LB1 with 2% NP-40 and 1% Triton X-100) for 20 min. Intact nuclei from this and the saved supernatant were collected by centrifugation at 1,700 x g for 5 min at 4°C, resuspended and washed in LB2 (10 mM Tris-HCl pH 8.0, 1 mM EDTA, 0.5 mM EGTA, 200 mM NaCl, protease cocktail inhibitor) for 10 min with rotation at 4°C, and collected again by centrifugation at 1,700 x g for 5 min at 4°C. Nuclei were gently rinsed 2x with LB3 (10 mM Tris-HCl pH 8.0, 1 mM EDTA, 0.5 mM EGTA, 0.01% NP-40, protease cocktail inhibitor) without disturbing the pellet. Nuclei were resuspended in 1 mL LB3 and transferred to a 1 ml milliTUBE (Covaris). Chromatin was sheared to a range of 300- 600 base-pair fragments using a Covaris E220 evolution Focused ultrasonicator with the following parameters: Peak power (140.0), Duty Factor (5.0), Cycles/Burst (200), Duration (600 seconds), Temperature (4°C). Sheared chromatin was quantified by Bradford Assay, and 450 µg of chromatin was brought to 500 µL in LB3 and 500 µL of ChIP Cocktail Mix (40 mM Tris-HCl pH 7.6, 150 mM NaCl, 1 mM EDTA pH 8.0, 1% Triton X-100, 0.5% NP-40, Protease inhibitor cocktail) was added. Prepared chromatin was pre-cleared by incubation with 20 µL of pre-washed Dynabeads Protein G magnetic beads (Invitrogen, 10004D) for 3 hours at 4°C with rotation. After bead removal, 10% input (100 µL) of pre-cleared chromatin was removed and set aside. Pre-cleared chromatin was immunoprecipitated with 5 µL of H3K27me3 antibody (Cell Signaling 9733) overnight at 4°C with constant rotation. Protein G magnetic beads (35 µL/IP) were blocked in buffer containing PBS, 0.5% BSA, and 20 µg Herring Sperm DNA (Sigma, D7290) with rotation at 4°C overnight. Blocked beads were washed 3X with PBS and 0.5% BSA and 2X with WB1 (50 mM Tris-HCl pH 7.6, 150 mM NaCl, 5 mM EDTA pH 8.0, 0.5% NP-40, 1% Triton X-100). Immuno-chromatin complexes were incubated with blocked beads for 3 hours with rotation at 4°C. Bead-immuno-chromatin complexes were then washed 3X for 5 min with rotation at 4°C with WB1, 3X with WB2 (50 mM Tris-HCl pH 7.6, 500 mM NaCl, 5 mM EDTA pH 8.0, 0.5% NP-40, 1% Triton X-100), 2X with WB1, and 1X with Low Salt TE (10 mM Tris-HCl pH 8.0, 1 mM EDTA pH 8.0, 50 mM NaCl). Beads were incubated in 50 µl of Elution Buffer (10 mM Tris-HCl pH 8.0, 10 mM EDTA, 150 mM NaCl, 5 mM DTT, 1% SDS) at 65°C for 15 min in 50 µL volume to elute immuno- chromatin complexes. The elution step was repeated, and eluates combined. Eluents and input were incubated overnight at 65°C with constant shaking to reverse crosslinks, followed by incubation at 37°C for 1 hour with DNase-free RNase A, then at 37°C for 2 hours with 10 µL of Proteinase K (20 mg/mL stock). DNA was isolated with a 1.5x ratio of KAPA Pure Beads (KAPA Biosystems KK8000) to DNA volume.

### siQ-ChIP library preparation and sequencing

Immunoprecipitated fragments and saved inputs were quantified with a Qubit dsDNA High Sensitivity Assay kit (Invitrogen Q32851), and 10 ng of purified DNA for each IP and input sample were used for library preparation with the KAPA Hyper Prep Kit (Kapa Biosystems KR0961). TAZ and Combo treatments required two IPs per biological replicate to attain enough material for library preparation, and this doubling has been accounted for in the parameters for siQ-ChIP (89,90) for these samples. Library preparation including fragment end-repair, A-tail extension, and adapter ligation was conducted per the manufacturer’s instructions (KAPA). Adapter-ligated fragments were amplified with 11 cycles following the recommended thermocycler program, and DNA was purified with two rounds of purification using KAPA Pure Beads (KK8000). Quality and quantity of the finished libraries were assessed using a combination of Agilent DNA High Sensitivity chip (Agilent Technologies, Inc.), QuantiFluor® dsDNA System (Promega Corp., Madison, WI, USA), and Kapa Illumina Library Quantification qPCR assays (Kapa Biosystems). Individually indexed libraries were pooled and 75 bp, paired end sequencing was performed on an Illumina NextSeq 500 sequencer using a 150 bp HO sequencing kit (v2) (Illumina Inc., San Diego, CA, USA) or 50 bp, paired end sequencing was performed on an Illumina NovaSeq6000 sequencer using an S2, 100 bp sequencing kit to a minimum read depth of 50M read pairs per IP library and 100M read pairs per Input library. Base calling was done by Illumina RTA3 and output of NCS was demultiplexed and converted to FastQ format with Illumina Bcl2fastq (v1.9.0).

### siQ-ChIP-seq processing and analysis

siQ-ChIP sequencing reads were 3’ trimmed and filtered for quality and adapter content using TrimGalore (v0.5.0) and quality was assessed by FastQC (v0.11.8). Reads were aligned to human assembly hg38 with bowtie2 (v2.3.5) and were deduplicated using removeDups from samblaster (v.0.1.24) (143). Aligned BAM files were used for quality control analysis with “deeptools” (v3.2.0) ‘plotFingerprint’ and ‘plotPCA’ functions. Aligned SAM files were then processed for pair-end reads with high mapping quality (MAQ >= 20), correct pair orientation (Sam Flags = 99, 163), and fragment length as described for siQ-ChIP (https://github.com/BradleyDickson/siQ-ChIP). Param.in files were prepared for each sample with all required parameters and measurements required for siQ-ChIP normalization. IP tracks (with H3K27me3 efficiency values) and comparative responses between drug treatments (relative to Vehicle) were generated with execution of getsiq.sh (version: February 2022) with the EXPlayout file (NOTE: params.in and EXPlayout file are provided with the GEO accession). Each individual inhibitor-treated biological replicate was compared to each individual vehicle-treated biological replicate.

To determine the change in H3K27me3 distributions between inhibitor-treated samples and vehicle-treated samples, each inhibitor-treated biological replicate (e.g. DAC301) was individually compared to the two vehicle-treated biological replicates (e.g. DAC301vsVeh1, DAC301vsVeh2), and the average log_2_ fold-change in response (area of peak in inhibitor-treatment/area of peak in vehicle treatment) was calculated. Next, conserved peaks between inhibitor-treated biological duplicates were determined by calculating the proximity of replicate 1 peaks to replicate 2 peaks (e.g. DAC301, DAC302) using the ‘closest’ command from “bedtools” (v2.25.0). Peaks were considered conserved among biological replicates if the peaks overlapped or were within 200 bp of each other. Finally, the average log_2_ fold-change in response was calculated for the peaks conserved between the two inhibitor-treated biological replicates. Peaks were considered significant if the log_2_ fold-change in response was ≥ 1.0 (increase in H3K27me3) or ≤ −1.0 (decrease in H3K27me3).

### ChromHMM (v1.23)

Custom ChromHMM (144) annotations for the HCT116 cell line were built with the following publicly available datasets:

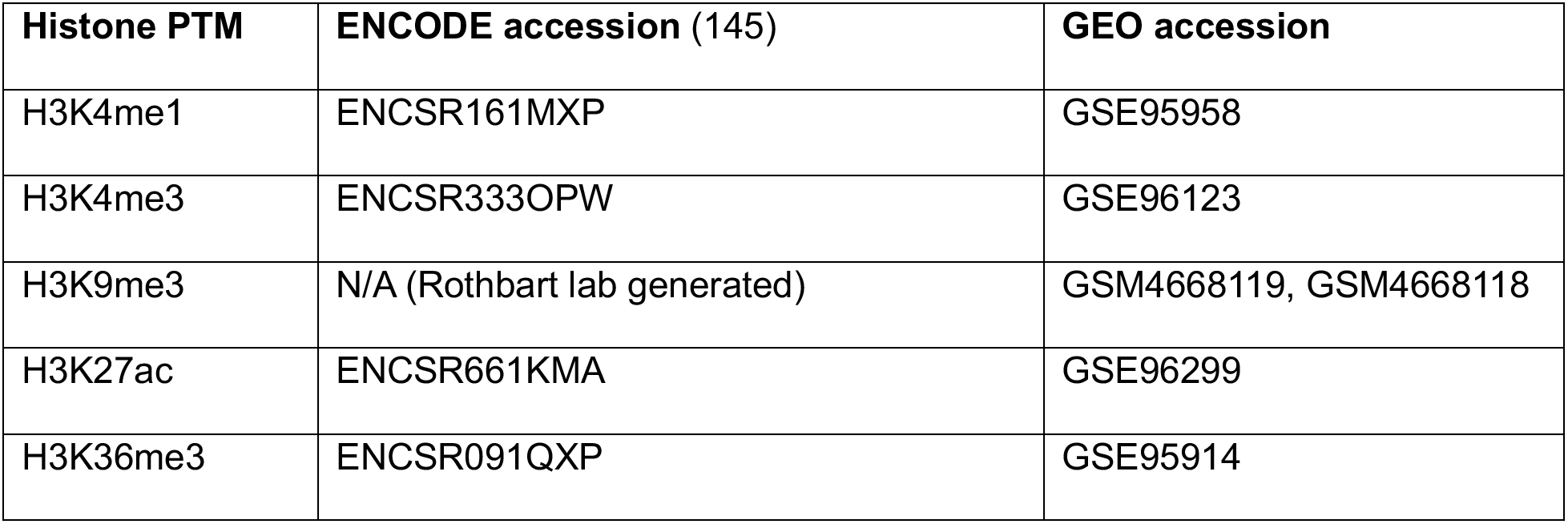

| Histone PTM | ENCODE accession (145) | GEO accession |
| --- | --- | --- |
| H3K4me1 | ENCSR161MXP | GSE95958 |
| H3K4me3 | ENCSR333OPW | GSE96123 |
| H3K9me3 | N/A (Rothbart lab generated) | GSM4668119, GSM4668118 |
| H3K27ac | ENCSR661KMA | GSE96299 |
| H3K36me3 | ENCSR091QXP | GSE95914 |

For H3K27me3 ChIP-seq dataset, we used our vehicle treated HCT116 H3K27me3 ChIP-seq data generated in this study. The original raw fastq.gz files were downloaded, re-processed, and aligned using the workflow described under siQ-ChIP processing and analysis. All aligned bam files were binarized using ‘BinarizeBam’ and then fed into ‘LearnModel’ to build a 15-state chromatin model. Chromatin states were assigned manually by considering the enrichment emissions for histone PTMs (**Figure S4H**), coverage of the genome (**Figure S4I**), and proximity to TSSs.

Enrichment overlap analysis with siQ-ChIP peaks, Repli-Seq phases, and differently expressed genes were conducted with the ‘OverlapEnrichment’ function using the genomic coordinates for each dataset.

### Repli-seq data accession and analysis

16-phase RepliSeq data (measuring replication timing from early to late replication) was downloaded (GEO: GSE137764 (93)) and each phase of replication timing was separated into their own genomic coordinates (both bed and bigwig files) for use in integrative ChromHMM, siQ-ChIP-seq, and gene expression analysis.

### Enzymatic Methyl (EM)-seq library preparation and sequencing

Libraries were prepared by the Van Andel Institute Genomics Core from an input of 41 ng to 51 ng of ChIP DNA (taken directly from DNA IP’d for siQ-ChIP-seq) using the NEBNext Enzymatic Methyl-seq Kit (New England Biolabs E7120L). The denaturation method used was 0.1 N sodium hydroxide, according to the protocol, and 8 cycles of PCR amplification were performed. Quality and quantity of the finished libraries were assessed using a combination of Agilent High Sensitivity DNA chip (Agilent Technologies 5067-4626) and QuantiFluor dsDNA System (Promega E2670). 150 bp paired-end sequencing was performed on an Illumina NovaSeq6000 sequencer using an S4, 300 bp sequencing kit (Illumina Inc., San Diego, CA, USA), with 10% PhiX to a minimum read depth of 100M read pairs per library. Base calling was done by Illumina RTA3 and output of NCS was demultiplexed and converted to FastQ format with Illumina Bcl2fastq (v1.9.0).

### EM-seq processing and analysis

EM-seq reads were aligned to human genome build hg38, duplicate marked (samblaster), and sorted (samtools) using the ‘biscuitBlaster’ pipeline from BISCUIT (v0.3.16) (https://huishenlab.github.io/biscuit/). Cytosine retention and callable SNP mutations were computed with ‘biscuit pileup’ and output into a VCF file. Cytosine beta values and coverage were extracted with ‘biscuit vcf2bed’ and CpG methylation status was merged with ‘biscuit mergecg’.

Quality control of the ChIP-EM-seq dataset was assessed several ways. First, PCA analysis (‘plotPCA’ deeptools(v3.2.0)) of both the siQ-ChIP and EM-seq libraries demonstrated that samples clustered based on their drug treatment (**Figure S4D**). To determine that EM-seq libraries truly were enriched for H3K27me3 fragments, k-means clustering from deeptools (n=4 clusters) was conducted to stratify protein-coding genes around the TSS by the efficiency of H3K27me3 IP (**Figure S4E**, top panel). Next, we calculated the average CpG read coverage in the EM-seq libraries for each of the H3K27me3 clusters and show that the clusters with the highest H3K27me3 level (C1,C2) also have the highest average read coverage, indicating that our approach effectively enriched for H3K27me3 in a quantitative manner and was sufficiently covered by EM-seq to interrogate DNA methylation levels (**Figure S4E**, bottom panel). CpH dinucleotides (control for EM conversion) showed less than 0.5% cytosine methylation indicating we achieved > 99% conversion efficiency on all EM-seq libraries (**Figure S4F**).

To assess the direct relationship between H3K27me3 and DNA methylation levels, we divided the genome into 100 bp bins, calculated the average H3K27me3 efficiency between biological replicates for each 100 bp bin, calculated z-scores for each bin considering the H3K27me3 efficiency across all bins in the vehicle treated samples, and finally subdivided the bins based on z-score into different H3K27me3 categories (**Figure S4G**). CpG coordinates were intersected with the 100 bp H3K27me3 bin coordinates using ‘intersect’ from bedtools (146) and calculated the average beta-value of the CpGs contained within each 100 bp H3K27me3 bin.

For integrative analysis with ChromHMM, siQ-ChIP, and gene expression, cytosine retention and coverage values were combined and a CpG was retained for downstream analysis if it was covered by ≥ 8 sequencing reads. Bed files were constructed for each sample with the calculated beta-value for each CpG and converted to bigwigs using UCSC Browser Tools (v March 2017).

### Integrative genomic analysis

As described in the respective sections above, bigwig files were generated for each sample for genome-wide H3K27me3 efficiency (siQ-ChIP-seq) and DNA methylation beta-values (EM-seq). Bed files with genomic coordinates for differential H3K27me3 peak analysis, ChromHMM chromatin state annotations, and Repli-seq phases were generated as described above. All data used for integrative analysis can be accessed under GEO accession# XXXXXXX or ENCODE (EZH2 ChIP-seq fold-change bigwigs: ENCSR046HGP). All datasets were aligned to genome build hg38.

Integrated siQ-ChIP-seq and EM-seq analysis was conducted with deeptools (v3.2.0) (147) by constructing matrices with ‘computeMatrix’ across queried genomic coordinates with the respective bigwig data and visualizing the summarized integration with ‘plotProfile’ and ‘plotHeatmap’.

Bivalent Enhancers that demonstrated an increase in H3K27me3 efficiency following DAC treatment were identified by intersecting the peak coordinates with the chromHMM EnhBiv coordinates using ‘intersect’ from bedtools (146). Motif enrichment analysis was conducted on the intersected Bivalent Enhancer coordinates (-/+ 10kb) using ‘findMotifsGenome.pl-len 8,10,12’ from HOMER (v4.11.1) (148).

### Genomic DNA isolation

Genomic DNA was extracted using the DNeasy Blood & Tissue Kit (Qiagen 69504) following the standard protocol. Samples were then treated with 1 mg/ml RNAse A at 37°C for 30 minutes. DNA was re-precipitated with 1/10 volume 3 M sodium acetate pH 4.8 and 2.5 volumes 100% ethanol and stored overnight at −20°C. Precipitated DNA was pelleted by centrifugation at 17,090 x g for 30 minutes at 4°C. The pelleted DNA was washed twice with 70% ethanol, allowed to dry for 15 minutes, and resuspended in nuclease-free water.

### Infinium MethylationEPIC BeadChip (EPIC array)

Genomic DNA was quantified by with the Qubit dsDNA High Sensitivity Assay kit (Invitrogen Q32851), and 1.5 µg of genomic DNA was submitted to the VAI Genomics Core for quality control analysis, bisulfite conversion, and DNA methylation quantification using the Infinium MethylationEPIC BeadChIP (Illumina) processed on an Illumina iScan system following the manufacturer’s standard protocol (149,150).

### EPIC array data processing and analysis

All analyses were conducted in the R statistical software (v4.1.2) (**R Core Team**). Raw IDAT files for each sample were processed using the Bioconductor package “SeSAMe” (version 1.8.12) for extraction of probe signal intensity values, normalization of probe signal intensity values, and calculation of β-values from the normalized probe signal intensity values (151–153). The β-value is the measure of DNA methylation for each individual CpG probe, where a minimum value of 0 indicates a fully unmethylated CpG and a maximum value of 1 indicates a fully methylated CpG in the population. CpG probes with a detection p-value > 0.05 in any one sample were excluded from the analysis. Differentially methylated regions (DMRs) were called using the Bioconductor package “DMRcate” (v3.13), and regions were considered differentially methylated if at least five contiguous CpGs demonstrated a mean difference of 0.15 methylation change in the drug treated cells compared to vehicle treated HCT116 cells.

### Construction and Sequencing of Directional total RNA-seq Libraries

Libraries were prepared by the Van Andel Institute Genomics Core from 500 ng of total RNA using the KAPA RNA HyperPrep Kit (Kapa Biosystems, Wilmington, MA USA). Ribosomal RNA material was reduced using the QIAseq FastSelect –rRNA HMR Kit (Qiagen, Germantown, MD, USA). RNA was sheared to 300-400 bp. Prior to PCR amplification, cDNA fragments were ligated to IDT for Illumina TruSeq UD Indexed adapters (Illumina Inc, San Diego CA, USA). Quality and quantity of the finished libraries were assessed using a combination of Agilent DNA High Sensitivity chip (Agilent Technologies, Inc.) and QuantiFluor® dsDNA System (Promega Corp., Madison, WI, USA). Individually indexed libraries were pooled and 50 bp, paired end sequencing was performed on an Illumina NovaSeq6000 sequencer to an average depth of 50M raw paired-reads per transcriptome. Base calling was done by Illumina RTA3 and output of NCS was demultiplexed and converted to FastQ format with Illumina Bcl2fastq (v1.9.0).

### RNA-seq processing and analysis

Raw 50 bp paired-end reads were trimmed with TrimGalore! (http://www.bioinformatics.babraham.ac.uk/projects/trim_galore/) followed by quality control analysis with FastQC. Trimmed reads were aligned to GRCh38.p12 and indexed to GENCODE v29 via STAR (v2.5.3a) aligner with flags ‘-twopassMode Basic\-quantMode GeneCounts’ for feature counting.

ReadsPerGene output count files were constructed into a raw read count matrix in R. Low count genes were filtered (1 count in at least one sample) prior to edgeR (v3.36.0) count normalization and differential expression analysis with voomWithQualityWeights and quasi-likelihood fit set to robust. Principal component analysis was calculated using ‘prcomp’ in the R stats package on the normalized expression matrix. Differential expression analysis was conducted with voomWithQualityWeights and a quasi-likelihood fit set to robust. Each dataset was treated separately for differential expression analysis (i.e. Day 3 acute treatment, Day 6 prolonged treatment, and CsA treatments), and all inhibitor treatments were compared to their respective Vehicle samples. Genes were considered differentially expressed if |Log_2_FC| ≥ 1 and FDR ≤ 0.01. Venn diagrams of overlapping genes were generated using BioVenn (154).

### Transposable Element (TE) analysis

TE analysis of total RNA-seq datasets was conducted with SQuIRE (v0.9.9.92) (https://github.com/wyang17/SQuIRE) (155). TEs were aligned to hg38, counted, and analyzed for differential expression relative to the Vehicle-treated samples using ‘Map’, ‘Count’, and ‘Call’ commands, respectively. TEs were considered differentially expressed if |Log_2_FC| ≥ 1 and adjusted p- value ≤ 0.01.

### Gene Set Enrichment Analysis (GSEA)

GSEA (v4.1.0) (156,157) was conducted across the following curated gene set databases: HALLMARK, c2.cgp, c2.cp.biocarta, c2.cp.kegg, c2.cp.reactome, c3.tft.v2023.1. Phenotype comparisons were set to Combo VS REST for all analysis with weighted enrichment statistic and Signal2Noise settings for ranking genes. Maximum and minimum size of a gene set was set to 2500 and 15 genes, respectively. Genes marked as a “core enrichment” gene were used for heatmap row z-score analysis of normalized cpm values.

### TCGA analysis

‘Gene Expression Quantification’ data for ‘Transcriptome Profiling’ of primary solid tumors were queried, downloaded, and prepared using the Bioconductor package “TCGAbiolinks” (158) for the following datasets: Colorectal Adenocarcinoma (COAD READ (105)), Breast cancer (BRCA (106)), and Ovarian cancer (OV (107)). Prepared gene counts were normalized by library size and overall transcription level with the Bioconductor package ‘edgeR’. For each cancer dataset, the respective AZA IMmune gene set (AIM) derived from Li et al. (104) was used for supervised clustering (clustering_method = “average”; clustering_distance = “correlation”) of the patient normalized gene counts using pheatmap. Patients were assigned into AIM “hot” and “cold” clusters based on the first column dendrogram branch. After AIM categories were assigned for each patient, supervised clustering of the Core Enrichment genes from the KEGG Calcium Signaling gene set (identified from GSEA analysis of Day 3 acute treatment Combo VS REST) was done using pheatmap. For direct comparison of the AIM and Calcium Signaling gene sets, the average z-score across the respective gene set was calculated for each patient and used for Pearson correlation analysis.

### PRC2 Inhibition In Vitro Assay

Reactions (10 μL) containing 150 nM of PRC2 (comprising EZH2, SUZ12, EED and RbAp46/48; Active Motif, 31887), 1 µM recombinant nucleosomes wrapped with 187bp DNA (EpiCypher 16-2004), and 1 μCi of ^3^H-SAM (PerkinElmer) in KMT reaction buffer (50 mM Tris pH 8.8, 2 mM MgCl_2_, 0.02% Triton X-100, and 1 mM dithiothreitol) with indicated amounts of EZH2 inhibitors or an equivalent concentration of DMSO were incubated for 1 hour at room temperature. Reactions were stopped by addition of trifluoracetic acid to a final concentration of 0.5%, neutralized by diluting with 150 μL of 50 mM NaHCO_3_, and transferred to streptavidin-coated FlashPlates (PerkinElmer). Plates were incubated for 15 min, sealed, and counted in a MicroBeta2 liquid scintillation counter (PerkinElmer MicroBeta2) for 1 min per sample. Percent activity was calculated by comparing to DMSO control and IC_50_ values were calculated using GraphPad Prism.

