## Supplemental Figures for "Select EZH2 inhibitors enhance the viral mimicry effects of DNMT inhibition through a mechanism involving calcium-calcineurin-NFAT signaling"

### Fig. S1 Chomiak et al.

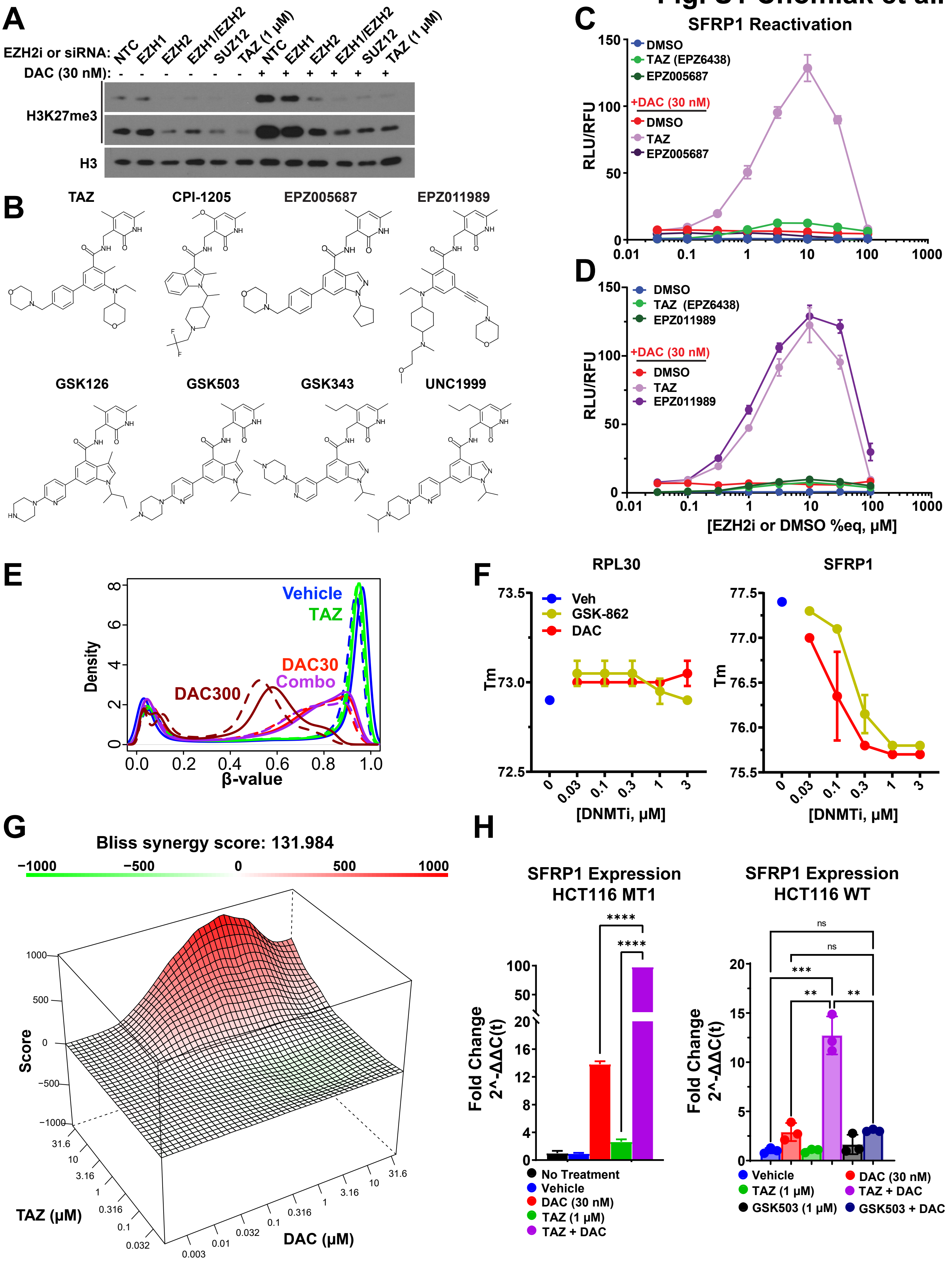

Fig. S2 Chomiak et al.

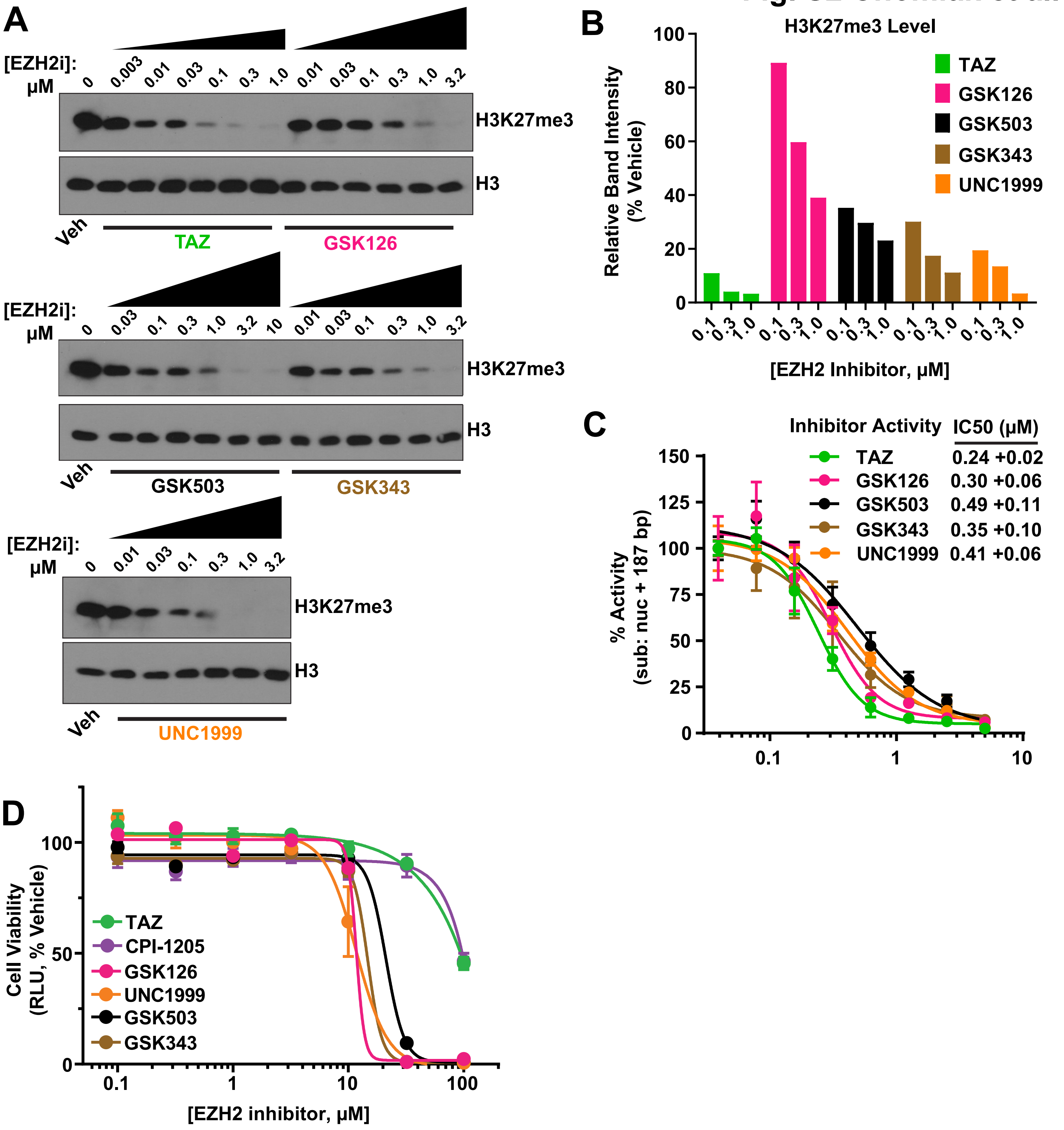

Fig. S3 Chomiak et al.

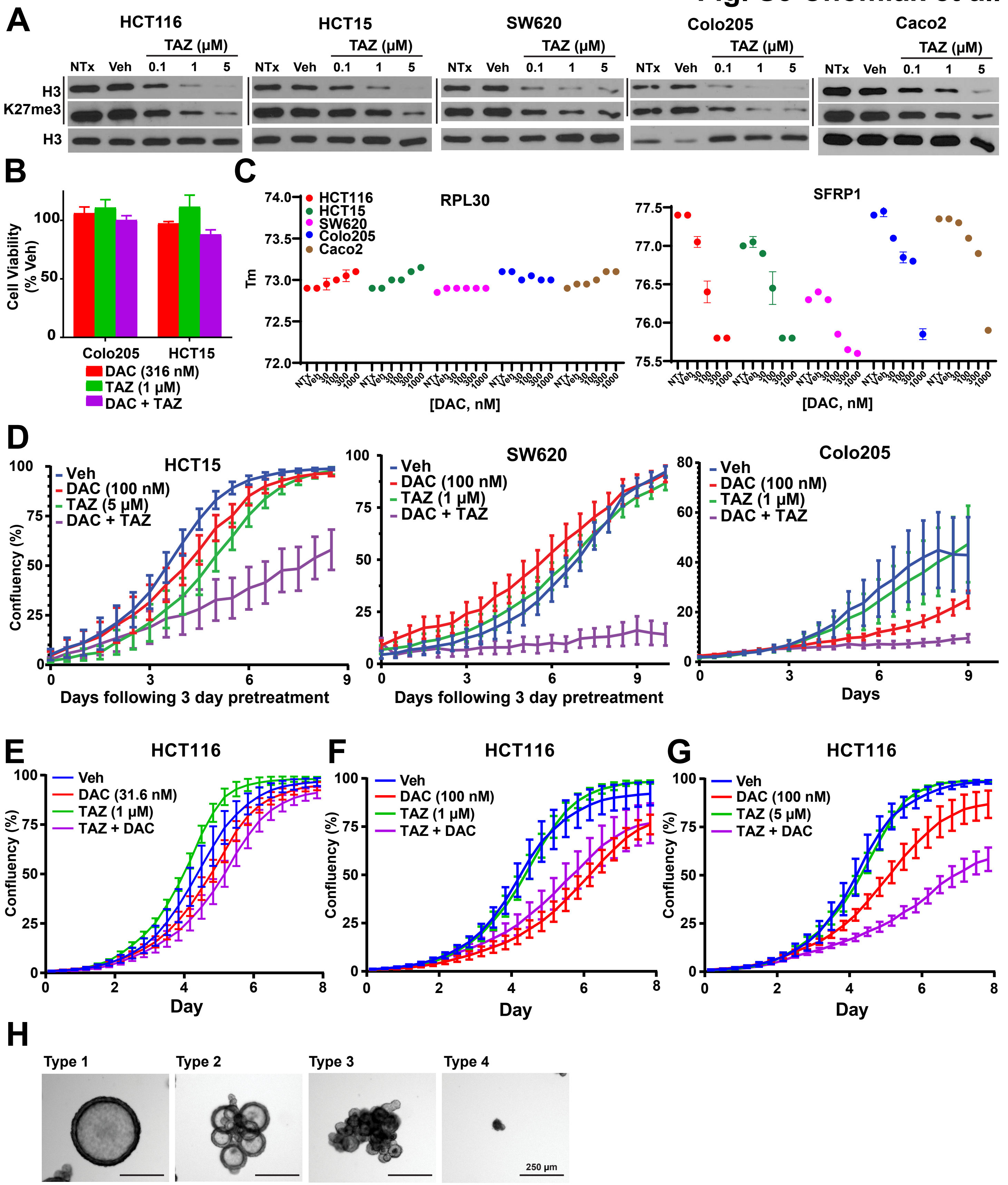

### Fig. S4 Chomiak et al.

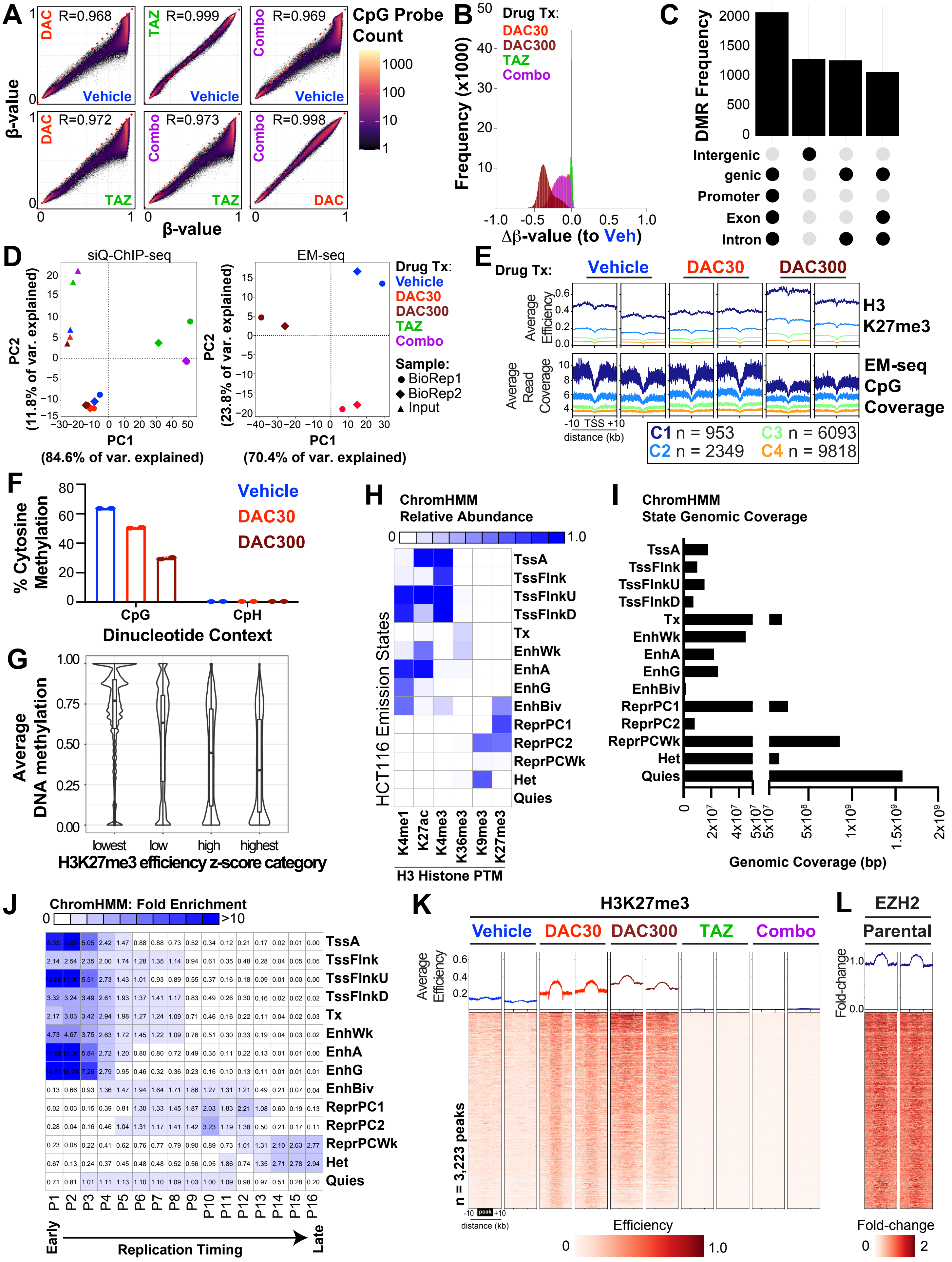

### Fig. S5 Chomiak et al.

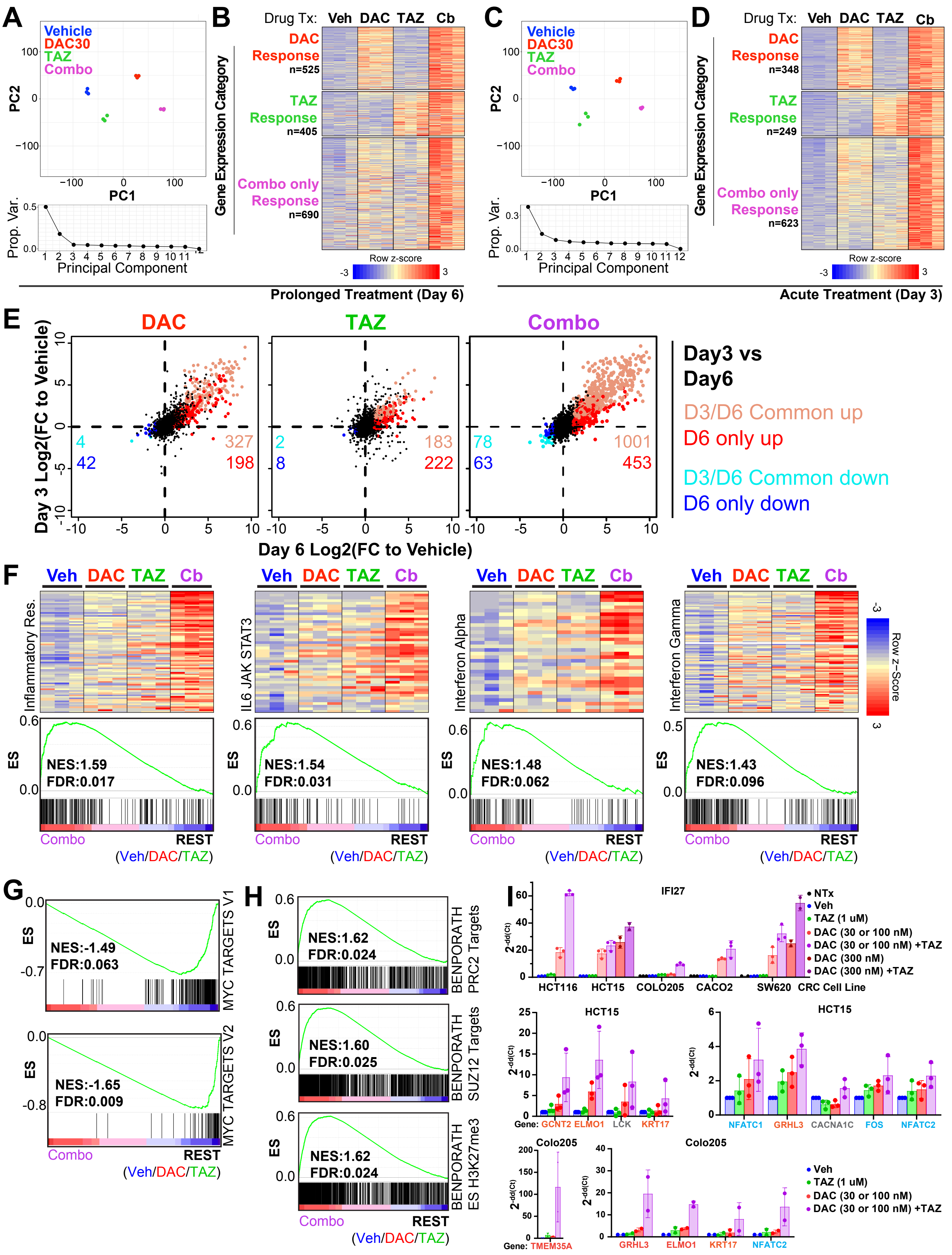

#### Fig. S6 Chomiak et al.

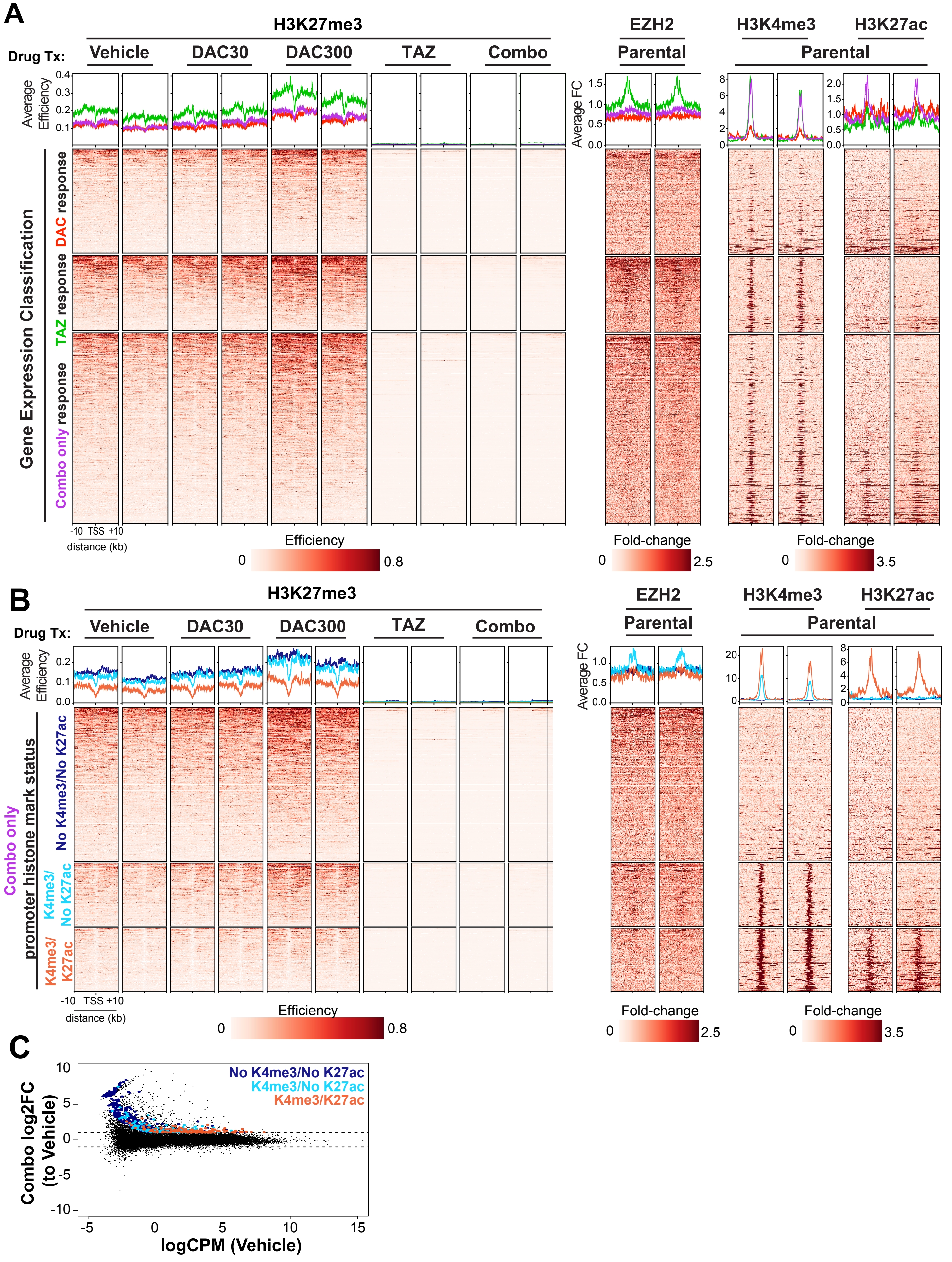

**Fig. S7 Chomiak et al.**

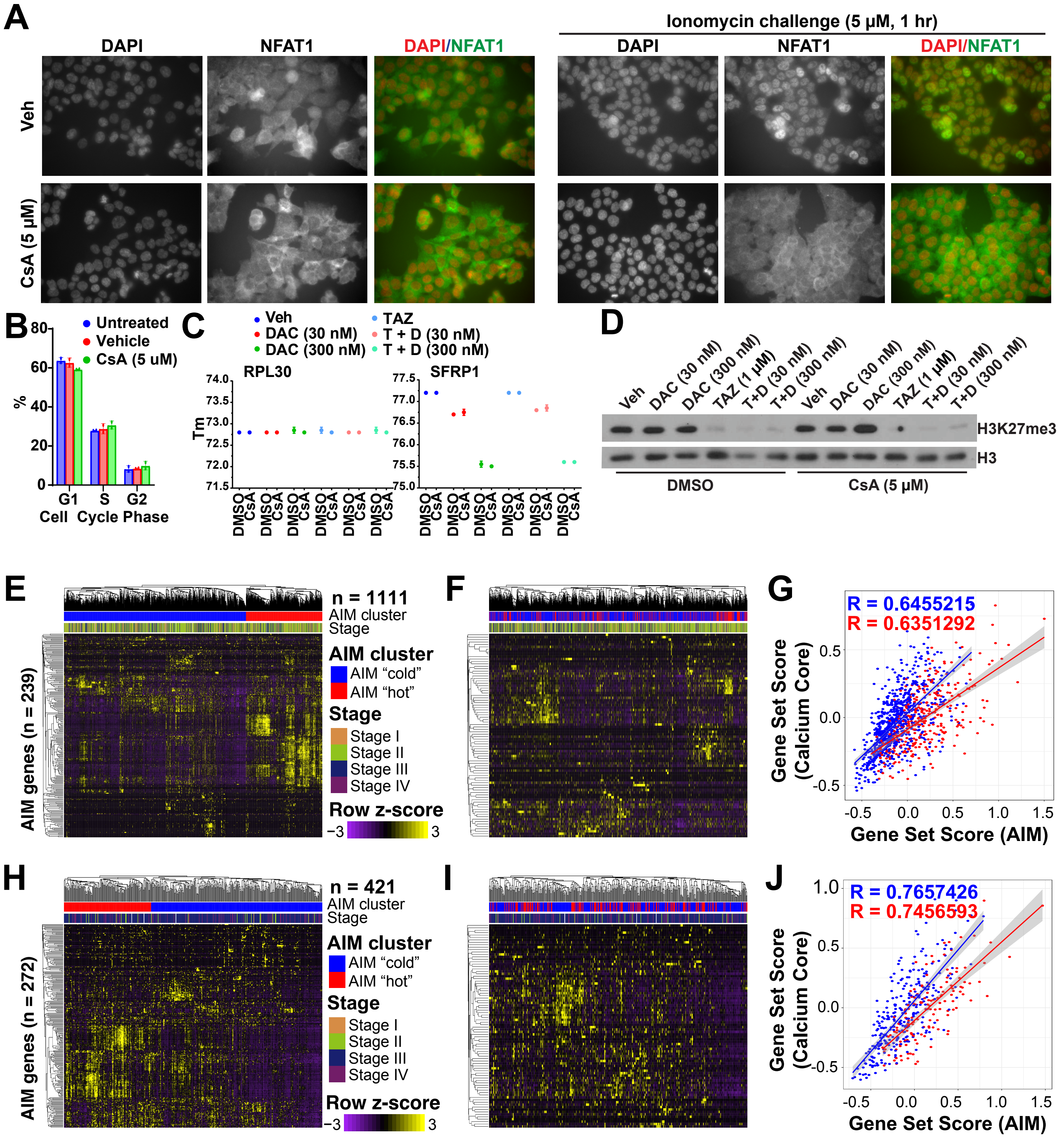
